# Sources of path integration error in young and aging humans

**DOI:** 10.1101/466870

**Authors:** Matthias Stangl, Ingmar Kanitscheider, Martin Riemer, Ila Fiete, Thomas Wolbers

## Abstract

Path integration is a vital function in navigation: it enables the continuous tracking of one’s position in space by integrating self-motion cues. Path integration abilities vary across individuals but tend to deteriorate in old age. The specific causes of path integration errors, however, remain poorly characterized. Here, we combined tests of path integration performance with a novel analysis based on the Langevin diffusion equation, which allowed us to decompose errors into distinct causes that can corrupt path integration computations. Across age groups, the dominant errors were due to noise and a bias in speed estimation. Noise-driven errors accumulated with travel distance not elapsed time, suggesting that the dominant noise originates in the velocity input rather than within the integrator. Age-related declines were traced primarily to a growth in this unbiased noise. Together, these findings shed light on the contributors to path integration error and the mechanisms underlying age-related navigational deficits.

## Introduction

Spatial navigation is a complex behavior that combines many computations, including the storage and recall of information, the integration of information from multiple sensory and non-sensory brain areas, planning, prediction, and decision making. A vital component of navigation-related computations is path integration – the integration over time of a self-motion estimate, in the strict sense of vector calculus, to maintain an updated estimate of one’s position and orientation while moving through space.

Self-motion estimates themselves derive from a sophisticated pooling over multiple sensory modalities, and rely on proprioceptive and vestibular information, visual optic flow signals (i.e., the pattern of apparent motion of objects, surfaces and edges), as well as motor efference copies that are produced during movement (Etienne & Jeffery, 2004). After being processed in their respective low-level sensory systems, these cues are integrated in brainstem nuclei as well as cortical structures (area MST) to allow an overall estimation of angular and linear movement velocity (Bassett & Taube, 2001; Biazoli et al., 2006; Angelaki & Cullen, 2008; Britten, 2008; Clark et al., 2012; Cullen, 2012; Butler & Taube, 2015). The integration of these cues is an error-prone process, and previous studies have demonstrated that path integration abilities therefore vary largely across individuals (Loomis et al., 1993; Klatzky et al., 1999; Chrastil et al., 2017). However, we have only a limited understanding of the specific sources of error that may corrupt path integration computations. In this work, we obtain quantitative measurements of path integration performance in subjects of different ages and develop and apply a method to decompose the observed path integration errors into components that can shed light on the mechanisms that underlie the observed errors (cf. Brunton et al., 2013).

A circuit that functions as a path integrator for two-dimensional space must do the following: take as input the given two-dimensional velocity signal; remember the previous integrated state; increment the previous integrated state by adding to it a quantity proportional to the instantaneous velocity input. There are thus several natural sources of error: First, the velocity estimate might be wrong, with systematic bias or unbiased noise. Second, the integrator might remember its past states in a leaky way, so that there is a decay of information over time. Third, the velocity input-based increments might be summed with a scaling or gain prefactor that differs from the value required to match the instantaneous displacement. Fourth, the integrator might itself be noisy.

These errors accrue over the course of a spatial movement trajectory, and the net localization error at path’s end will depend on the details of the trajectory. Thus, properly modeling and decomposing these errors requires iteration of a temporal dynamics, a statistical model that incorporates these dynamics, and sufficiently rich and varied spatial trajectories in the input data. One final error arises when a downstream neural circuit or the human experimenter attempt to obtain a readout or report of the internal state of the integrator.

Our goal in the present work is not only to understand the contributors to path integration error, but also to reveal sources of age-related degradation in navigation performance. Specifically, aging has deleterious effects on path integration ability, with declines in the triangle completion task (Loomis et al., 1993) – a standard assay of path integration performance (Allen et al., 2004; Mahmood et al., 2009; Adamo et al., 2012; Harris & Wolbers, 2012). Older adults are less accurate in reproducing travel distances or rotations (Mahmood et al., 2009; Adamo et al., 2012; Harris & Wolbers, 2012), and they exhibit worse path integration performance even if additional landmark information is available (Harris & Wolbers, 2012; Bates & Wolbers, 2014). Despite the sizeable body of research on losses in path integration performance with age, little is known about which specific aspects of the path integration computation or process are most affected in old age.

To address this important question, we combine an immersive virtual reality path integration experiment with a novel mathematical approach to reveal the sources of path integration error. We characterize the different contributors to error across subjects, and study group differences between young and older adults.

## Results

Young and older adults experienced a virtual reality environment from a first-person perspective via a head-mounted display (HMD). When participants moved in the real world, their movements were translated into movements (i.e., changes in location and viewing orientation) in the virtual environment, allowing them to walk around within the virtual world and use both body-based and visual self-motion cues to estimate their changing location.

For the path integration task, subjects were asked to track their own position and orientation as they were guided through this environment along pre-defined but unmarked curved paths (**Figure 1**). Each path had four intermediate stopping points, at which participants were asked to stop and report their estimate of the direct distance and direction to the path’s starting point.

**Figure 1:**
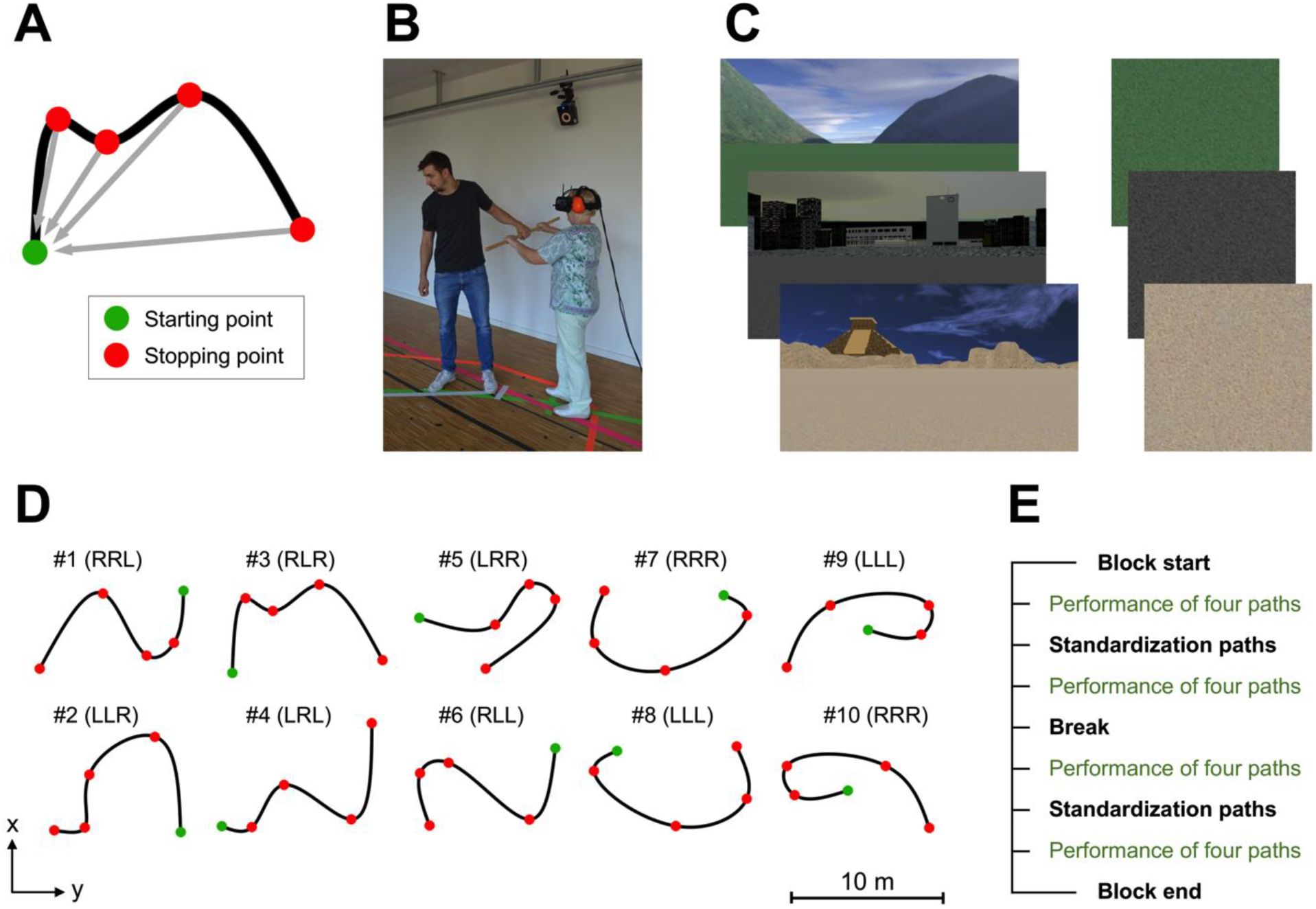
Path integration task. **A:** Example path, viewed from a top-down perspective. Participants began at the starting point (green dot) and then walked along the path (curved black line). There were four stopping points (red dots) along each path; at these points, participants were asked to report their estimate of the direct distance and angle to the path’s starting point (gray arrows). **B:** During the experiment, participants saw a virtual environment from first-person perspective via a head-mounted display (HMD). All movements in the real world were tracked with a motion tracking system and translated to movements (i.e., changes in location and viewing orientation) in the virtual environment, so that participants could walk around and perform the task in the virtual environment. Participants held a wooden stick and were guided by the experimenter along a path. At each stopping point, the direct distance to the starting point had to be estimated verbally in meters and centimeters, and participants turned their body on the spot to indicate the orientation to the starting point. **C:** The three different virtual environments (left panel) used in the path integration task. Each environment comprised a ground plane and distant landmark cues. As these landmark cues were rendered at infinity, they can facilitate the determination of heading direction, but do not allow participants to determine their position or distance information. One tile of each environment’s ground plane is shown in the right panel. These tiles were textured to provide optic flow during movement, but were seamless (no visible border between adjacent tiles) and provided no fixed cues with positional information. Across environments, the ground planes differed only in color. **D:** Overview of the 10 different paths used in the experiment. Each path contained three turns, and turn directions (i.e., left “L” and right “R” turns) were counter-balanced between paths. **E:** Each participant performed three blocks of the path integration task. Each block consisted of 16 paths (paths #1-10, and paths #1-6 repeated without intermediate stopping at stopping points 1-3) in pseudo-randomized order. In addition, after the 4th and 12th path of each block, participants performed so-called “standardization-paths” (i.e., straight lines with a length of 2m, 6m, and 10m), which were used during data analysis to correct for each participant’s abilities in verbally reporting distances using meter/centimeter units. Finally, there were short breaks between blocks and in the middle of each block. See Methods for more details.

Most participants showed a characteristic increase in path integration error over the course of their trajectories (**Figure 2A**). We first pooled path integration errors across individuals, separately for the group of young and older adults, and evaluated whether participants’ performance in the path integration task was better than random guessing. Indeed, estimates of location were highly correlated with true location (**Figure 2B**; r = 0.64 to 0.94, all p < 0.00001) while shuffled responses across trials (corresponding to different trajectories) per stopping point exhibited much larger squared errors (**Figure 2C**).

**Figure 2:**
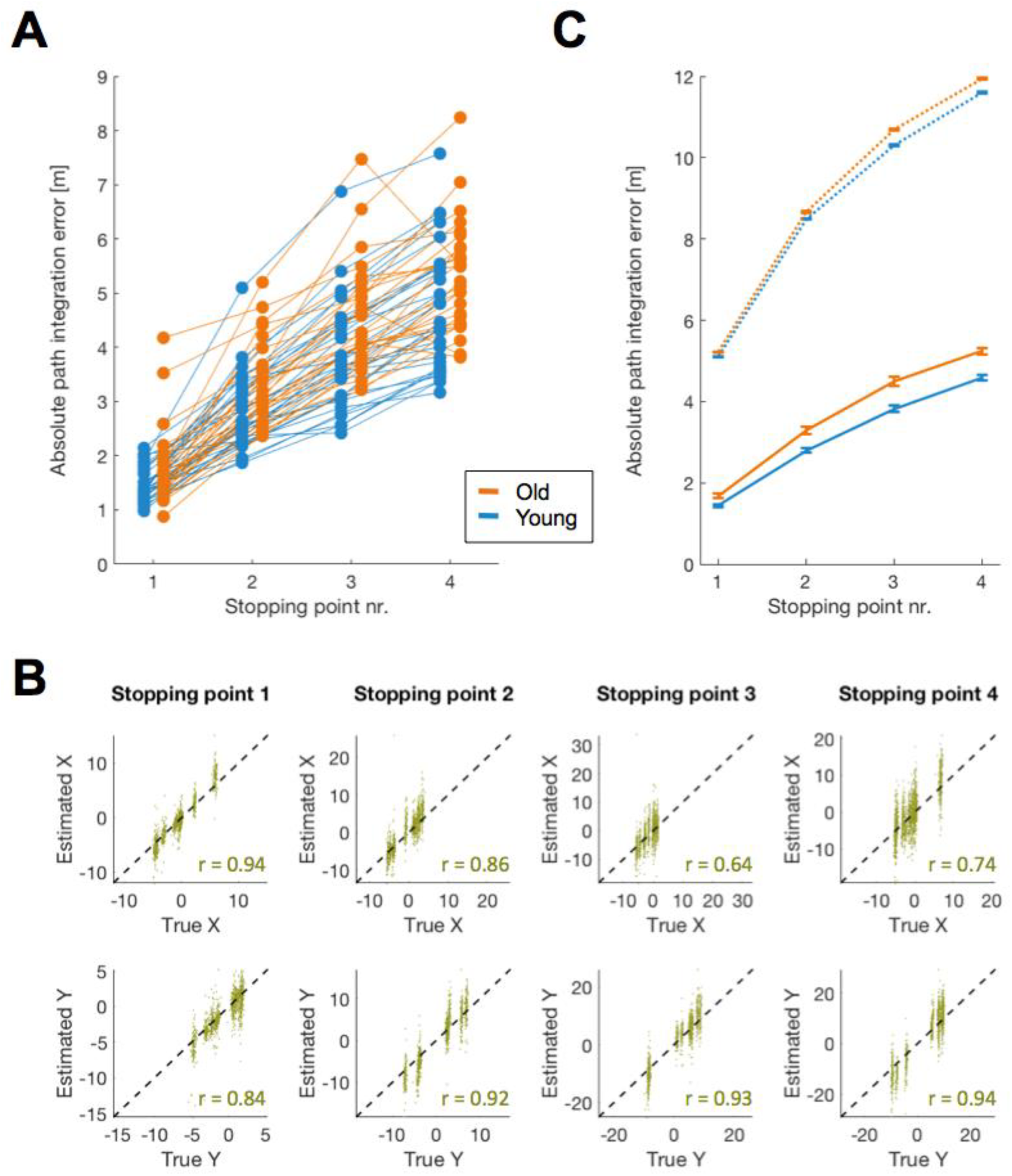
Path integration performance across both age groups. **A:** Absolute path integration errors over four stopping points for young and older adults. Average errors per stopping point are shown for each participant separately by blue (young adults) and orange (older adults) dots, connected with lines between stopping points. **B:** Each participant’s location estimate (y-axis) versus their true location (x-axis) at each of the four stopping points (columns), separately for x-coordinates (top row) and y-coordinates (bottom row). Plots show data from all participants and all paths. The diagonal (dashed line) indicates perfect response (estimated location = true location). All correlation coefficients are statistically significant (all p < 0.00001). All units are meters. **C:** Absolute path integration errors of young and older adults versus errors with shuffled responses. It is evident that the mean absolute path integration error of both groups (solid lines) is much lower than the errors obtained from shuffling each participant’s responses across trials. Error bars indicate group mean ±SEM.

### Dynamical model of errors

Next, we built and fit a temporally-resolved computational model of participants’ responses to disentangle different sources of path integration error. Path integration was modeled as a continuous updating of an internal location estimate by an integrator receiving a time-varying velocity estimate. The process was assumed to be corrupted over time by the following error sources: under- or overestimation of velocity (velocity gain bias), leaky integration of the velocity signal (memory decay or leak), an additive location bias (additive bias, AB), and ongoing zero-mean Gaussian additive noise (accumulating noise, AN), which accumulates and could be interpreted as originating in either the velocity input to the integrator or within the integrator. In our default “full model”, the accumulating noise is naturally interpreted as driven by the velocity input, as it accumulates during the trajectory and in proportion to travel distance, but does not accumulate during stopping points. In an alternative formulation that we tested, the noise accumulates over time instead of travel distance (i.e. it also accumulates during stopping points), and thus would be more naturally interpreted as internal to the integrator (as described in more detail below). In addition, we assumed that the participants’ reports of distance and angle to the starting point are imperfect and corrupted by reporting noise (RN), with angular and radial components.

Model parameters per participant were obtained by the best fit across all paths and trials (**Supplementary Figure S1**). The model captured not only the magnitude of errors averaged across paths (**Figure 3A**), but also predicted with high accuracy the full, time-resolved, signed errors at different portions of the several individual paths (**Figure 3B**).

**Figure 3:**
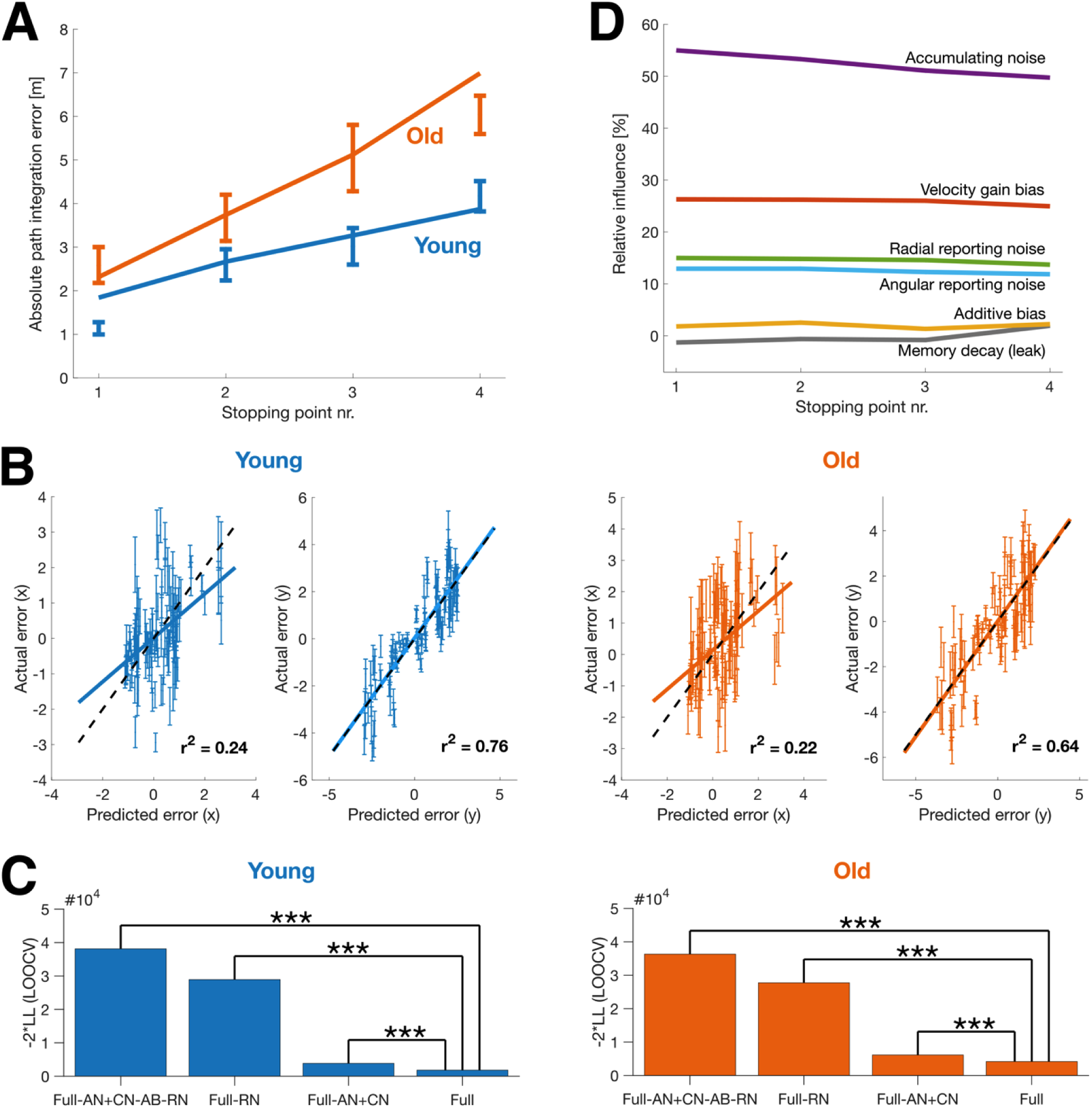
Computational modeling results. **A:** Path integration errors, averaged across all trials, of one example participant from each of the two age groups (error bars), and model fits (solid lines). Error bars represent variability (SEM) over trials. Data and fits for all individual participants and average fits by age group are in **Supplementary Figure S2**. **B:** Measured single-trial path integration error vectors versus error vectors predicted by the model. Predicted position is computed individually per subject per trajectory; datapoints show the per-trajectory predicted position, averaged across subjects of the same age group on the same trajectory and trial (to reduce scatter). Error bars represent the variability (SEM) at a single trial across participants (trial order was identical across participants). The dashed black (diagonal) line indicates perfect prediction; solid lines represent the best-fitting linear regression fit of the datapoints. All units are meters. **C:** Model comparison: negative log-likelihood scores using LOOCV between models, with higher bars indicating a poorer model fit. *** denotes “very strong” evidence against the model relative to the full model (ΔBIC (ΔLOOCV)> ≫10; see also Methods section and **Supplementary Figure S3**). Key to model names: The full model (“Full”) is our default, with ongoing “accumulating noise” (AN) that is proportional to the length of the travelled path, nonzero additive bias (AB) and velocity gain bias parameters, and reporting noise (RN) that is proportional to the magnitude of the reported variable. CN refers to when the non-reporting portion of the noise is constant rather than accumulating. CRN refers to when the reporting noise is constant (rather than proportional to the magnitude of the reported variable). +/- refers to the addition/removal of that contribution to the model, respectively. **D:** Impact of different model parameters on the predicted path integration error. Relative influence measures the predicted reduction in square error by setting a parameter to its ideal value corresponding to noiseless and unbiased integration. Note that due to the nonlinearity of the model, the relative influences do not have to sum to 100%, and that a parameter’s relative influence can be negative if the reduced square error is larger than the square error of the full model (see Methods section for more details). Ongoing noise, whose effect is cumulative and proportional to travelled distance, has the largest relative influence.

We then quantified the support for the detailed structure of the full model by comparing it to other variants with fewer parameters or different noise models (i.e., ongoing noise that accumulated over time instead of travel distance, or a noise that remained constant instead of accumulating). We considered reporting noise that was proportional in magnitude to the reported variable, or constant in magnitude, or absent (**Figure 3C**). Model comparisons were carried out using both Bayesian Information Criterion (BIC) and leave-one-out cross-validation (LOOCV), which penalize overly rich models that do not improve prediction performance (see Methods section for more details about different model variants and BIC/LOOCV model comparisons).

The full model was highly favored (“very strong” evidence in support, or Δ*BIC* (Δ*LOOCV*) ≫ 10) relative to alternatives, including models with no reporting noise (Full-AN+CN-AB-RN, Full-RN) or non-accumulating (constant) noise (Full-AN+CN-AB-RN, Full-AN+CN), consistently across both age groups (Young: Full vs. Full-AN+CN-AB-RN, Δ*BIC* = 36303, ΔLOOCV = 34743; Full vs. Full-RN, Δ*BIC* = 27103, ΔLOOCV = 26089; Full vs. Full-AN+CN: Δ*BIC* = 2035, ΔLOOCV = 2021; Old: Full vs. Full-AN+CN-AB-RN, Δ*BIC* = 30731, ΔLOOCV = 32124; Full vs. Full-RN, Δ*BIC* = 22579, ΔLOOCV = 23577; Full vs. Full-AN+CN: Δ*BIC* = 1963, ΔLOOCV = 1957). Specifically, the full model outcompeted the Full-AN+CN-AB-RN variant, which – with non-accumulating noise, no additive bias in integration, no reporting noise, but biased and leaky velocity integration – is the closest analogue to a leading existing model of human path integration performance (Lappe et al., 2007, 2011).

The full model was also much better supported than the alternatives when parameters were fit individually for each participant, even after accounting for the much larger number of parameters than fitting a common set of model parameters by age group (**Supplementary Figure S4**; Young: Δ*BIC* = 7412, Old: Δ*BIC* = 5367). However, the relative preference for an additive bias in the integrator was inconclusive, and depended on both the comparison method (BIC and LOOCV) and subject group (**Supplementary Figure S3**).

We next sought to quantify whether accumulating noise in the integrator is better explained by ongoing noise as a function of travel distance or elapsed time. In principle, the former would be a movement-dependent noise that is likely to arise from external velocity inputs to a neural integrator, while the latter is likely to arise within the integrator, due for instance to noise within the grid cell circuit in entorhinal cortex (Hafting et al., 2005; Burak & Fiete, 2009). We therefore compared the full model, which assumes the accumulated noise scales with traveled distance, with the “time model” variant that assumes a scaling with elapsed time, and found much stronger support for the full model across both subject groups (**Figure 4A** and Methods; Young: Δ*BIC* = 194, ΔLOOCV = 222; Old: Δ*BIC* = 525, Δ*LOOCV* = 533).

**Figure 4:**
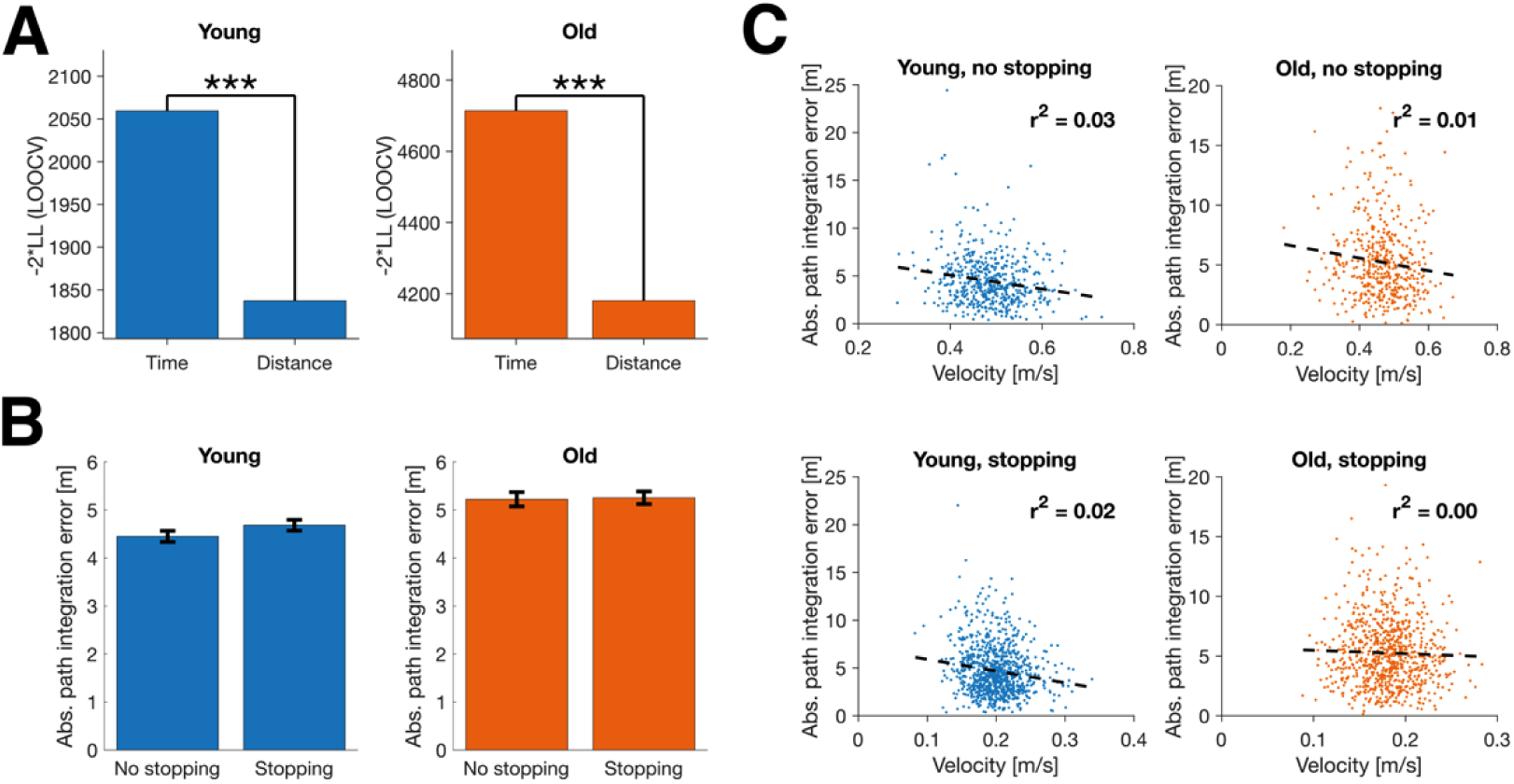
Time- vs distance-scaling of accumulating noise. **A:** Model comparison using LOOCV between the full model with accumulated error proportional to total travel distance versus total time in trajectory. For both age groups, the full model is better supported by the data. Higher bars indicate poorer model-fit. *** denotes “very strong” evidence against the model with poorer fit (ΔBIC (ΔLOOCV) ≫ 10; see Methods and **Supplementary Figure S5**). **B:** Average path integration error at the last stopping point, in trials with and without intermediate stopping points. The path integration error is very similar even though trials with stopping take much more time, indicating that the path integration error mainly scales with distance rather than time. Error bars indicate SEM. **C:** Walking velocity versus path integration error for trials with and without stopping and for both age groups.

More directly, we compared total error on trajectories in which subjects stopped versus did not stop at intermediate stopping points. Participants completed 48 paths in total, out of which 18 involved a stop only at the endpoint; in the remaining 30 paths, subjects also stopped at three intermediate stopping points to report the distance and angle to the starting point (see Methods section for more details). Since the different paths had very similar total length (17.7 ± 0.1 meters, mean ± SD), the total travel distances were similar over stopping and non-stopping trajectories, but the travel times differed substantially (88.7 ± 12.4 seconds versus 35.2 ± 3.9 seconds, mean ± SD). Nevertheless, path integration errors were very similar for stopping and non-stopping trajectories (**Figure 4B**), indicating that errors were mainly determined by the traveled distance instead of elapsed time, and therefore suggesting that the dominant source of accumulating noise is in the velocity inputs rather than within the integrator.

Moreover, given that paths had similar total lengths, the time-scaling model would predict a negative correlation between walking speed and path integration error: walking faster permits faster completion of the trajectory. However, we found little evidence for such negative correlation in the data (**Figure 4C**).

We next used the full model to assess the relative importance of the different sources of error during the task. To do so, we calculated the relative influence of each bias and noise parameter on the predicted square error (see Methods section for more details). We found the largest influence on total squared error to be from accumulating unbiased noise (50-55%) and the velocity gain bias (25-26%), followed by radial (14-15%) and angular (12-13%) reporting noise (**Figure 3D**). In contrast, the influence of both additive bias and memory leak were very small (< 3%), suggesting that the integrator itself is well-tuned to eliminate leak and internal bias and that the errors are due to velocity misestimation, with contributions from both an unbiased ongoing noise and a biased multiplicative gain in estimating speed.

Note that the result that the largest contribution to the error in the full model is from accumulating noise (**Figure 3D**) does not contradict the result that the introduction of reporting noise causes the largest increase in model fit (**Figure 3C**). Intuitively, **Figure 3C** can be interpreted as a measure of ‘error shape’, namely how different sources of error grow with traveled distance and distance to the starting point, while **Figure 3D** measures ‘error size’ in the context of the full model. In models without reporting noise, all errors have to be fit by a single noise source of incorrect shape, which causes the large discrepancy in **Figure 3C**.

### Age-related differences in path integration

Older adults performed less well in the path integration task compared to young adults. Absolute path integration errors were significantly higher in older adults by the first stopping point, and continued to be higher at all subsequent stopping points along the path (**Figure 5A**; stopping point #1: p = 0.016; #2: p = 0.004; #3: p = 0.005; #4: p = 0.005). Moreover, incremental path integration errors or the gain in error between adjacent stopping points (pooled over all stopping points; see Methods section for more details) were significantly higher for older relative to young adults (**Figure 5B**; p = 0.001).

**Figure 5:**
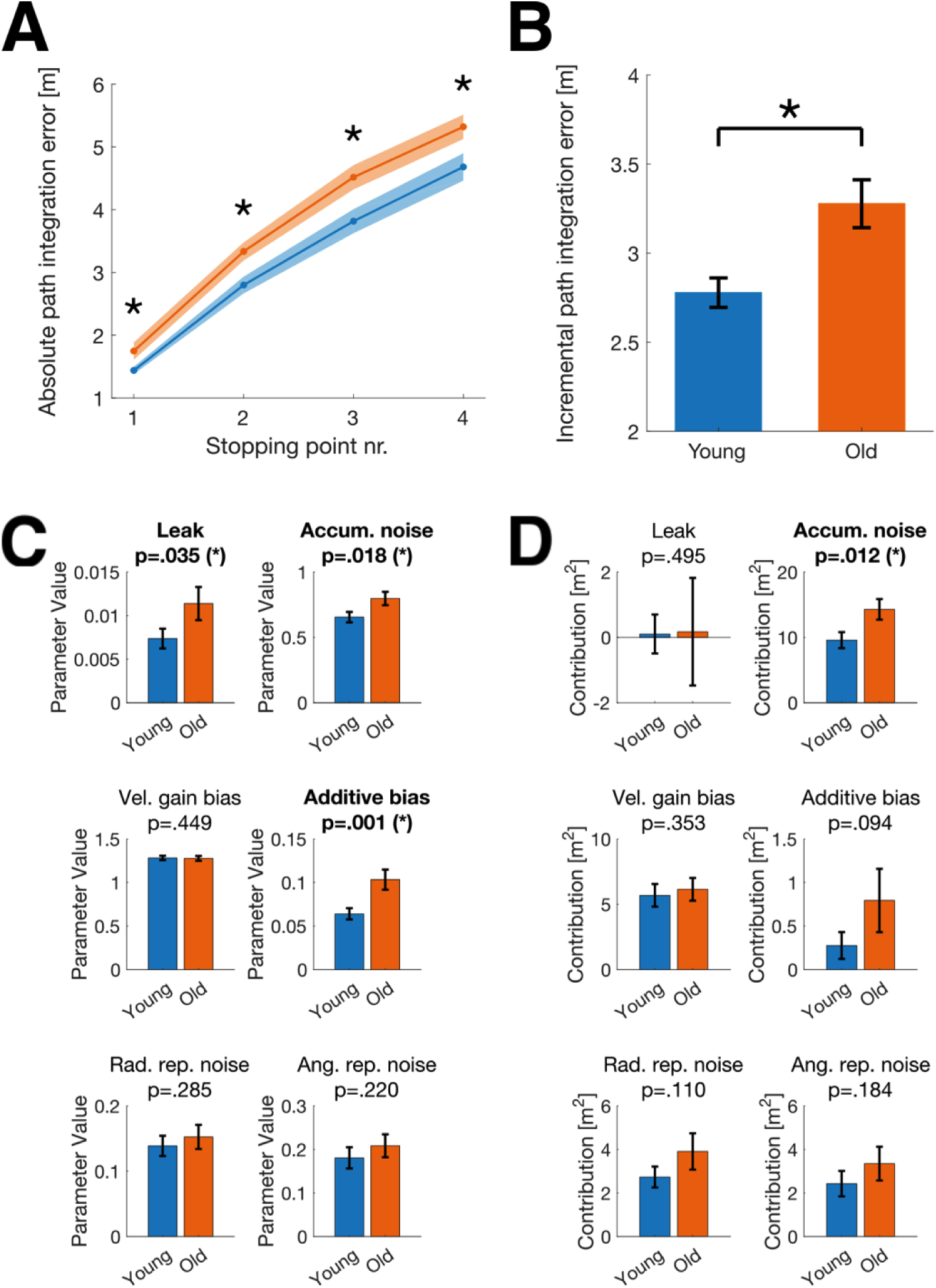
Path integration in older versus young adults. **A:** On average, older adults showed a higher absolute path integration error than young adults at all stopping points. Blue and orange lines indicate group mean ± SEM. **B:** The incremental path integration error (i.e., the additional contribution to the path integration error for each segment between adjacent stopping points), averaged across stopping points, was higher for older than young adults. **C:** Model parameter values, averaged over participants of the same age group. Parameter values for leak, accumulating unbiased noise, and additive bias were significantly higher in older relative to young adults. Individual parameter values for single subjects are shown in **Supplementary Figure S1**. **D:** Each model parameter’s contribution to the absolute square error, averaged over participants of the same age group. Only the accumulating unbiased noise resulted in a significant difference in error contribution between age groups. A parameter’s contribution is calculated by measuring the reduction in square error when setting the parameter to its ideal value corresponding to unbiased, noiseless integration (see Methods). Error bars indicate SEM. * denotes a significant difference between age groups (p < 0.05) in a one-sided permutation test with 10000 permutations.

To determine the underlying reason for the differential performance of older and young adults, we fit our computational model parameters individually across subjects, and then compared the extracted parameters between age groups. Older adults had a significantly larger additive bias (p = 0.001), a significantly larger amount of accumulating noise (p = 0.018), and greater memory leak (p = 0.035) than young adults (**Figure 5C**). However, some of these parameters had only a small overall contribution to the total error; comparing each parameter’s contribution to overall path integration error between age groups revealed that only the accumulating noise (p = 0.012) had a significantly higher contribution to error in older relative to young adults (**Figure 5D**), suggesting that velocity estimation degrades in relatively unbiased ways, to become noisier in older relative to young humans.

## Discussion

We used a novel immersive virtual reality path integration task in which young and older adults tracked their own pose (position and orientation) using visual and body-based motion cues while travelling along sinuous paths. Simultaneously, we developed a powerful analysis approach based on stochastic differential equations (the Langevin equation) to decompose path integration errors into temporally resolved gain, leak, bias, and noise terms and to estimate, on a trial-to-trial basis at different times along the path, how these different sources of error contribute to the location estimation error. In addition to sources of accumulating error, the analysis also included the possibility of errors in the generation of an explicit report of an internal estimate of the displacement vector, as subjects are asked to provide (at each stopping point). We performed mathematical inference of model parameters using an approach based on the Extended Kalman Filter. Disentangling the different sources of error allowed us to compare their influence on path integration errors across participants and between age groups.

A central assumption of our analysis is that observers track and use self-motion cues to continuously update their internal estimates of pose. However, path integration performance can also rely on a “configural strategy”, in which participants store the configuration of a path (i.e. segment lengths and turn angles) and only compute a homing response when required (Wiener et al., 2011). This strategy is often observed when the outbound path can be easily segmented into turns and distances – such as in the popular triangle completion task – and it can induce systematic biases such as a tendency to regularize turns and distances to canonical values (e.g., isosceles triangles or right-angle turns; Sadalla & Montello, 1989). To eliminate these confounds, we used irregularly shaped sinuous paths, in which translations and rotations were combined into curved trajectories. In addition, we asked subjects to report their internal estimates of the homing vector at intermediate stopping points. These strategies strongly encourage participants to continuously update their displacement estimates based on motion cues over the task.

Previous work by others including Lappe and colleagues (2007, 2011) examined distance misestimation when subjects indicated the magnitude of their displacement along straight or veering outbound paths, reporting that subjects systematically underestimate longer displacements. By contrast, we find that the dominant error in estimating two-dimensional displacement vectors comes from unbiased noise; systematic biases in leak and velocity gain contribute only modestly to total error. There are at least two ways in which the setups differ: First, the studies use different models to decompose error. All observed errors in Lappe et al. were projected onto two terms, the degree of leak in integration and the gain in velocity estimation. Our model included these terms but also allowed for accumulating noise and reporting errors, permitting richer possible interpretations of the possible contributors to the total error. To address whether the richer model is justified by the data, we performed Bayesian model comparison and cross-validation, and showed that the richer model exhibits better performance on unseen data than the simpler one. Thus, it is implausible that subjects exhibit substantial biases in velocity gain that are not discovered by the analysis model. Second, the subjects in Lappe et al. (2007, 2011) have access only to optic flow for motion estimation, while subjects in our study additionally have access to richer, body-based motion cues including vestibular signals. In rodent studies, when motion cues are less rich (passive transport on trolleys; head-fixed animals in virtual environments), displacements are underestimated, similar to the finding in Lappe et al., suggesting that a decreased availability of sensory motion cues in Lappe et al. may account for the dominant contribution of a velocity estimation bias in their findings.

Previous work (e.g., Petzschner & Glasauer, 2011; Petzschner et al., 2015) has considered the possibility that subjects learn and subsequently exploit information about regularities in the tasks they must perform. In a Bayesian interpretation, this information can be incorporated into prior assumptions or biases on the values that variables and parameters can take. Subjects have likely not performed the tasks we designed enough times to form useful priors to improve task performance over naive path integration, and the tasks have little repeatable or regular structure to be exploited. Nevertheless, the ability of our analysis method to isolate different sources of error and their impact on individual path integration performance can make it possible for future studies to investigate the existence or learning of biases, including ones related to a priori assumptions about the structure of the world.

Our discovery that path integration errors in (both young and old) human subjects are mainly explained by an unbiased noise – resulting in a random diffusion of the estimated locations away from their true values – suggests that both velocity estimation and integration are well-tuned to be fairly unbiased processes, i.e. that velocity is estimated with a gain near unity, and that integration is largely non-leaky. The unbiased noise must arise at some stage along the path integration process, and thus could in principle arise within the integrator (Compte et al., 2000; Brody et al., 2003; Boucheny et al., 2005; Wu et al., 2008; Burak & Fiete, 2009, 2012) or in the velocity inputs delivered to it. Given the large body of research suggesting that grid cells may be the neuronal substrates of path integration (Hafting et al., 2005; Fuhs & Touretzky, 2006; McNaughton et al., 2006; Burgess et al., 2007; Guanella et al., 2007; Hasselmo, 2008; Burak & Fiete, 2009; Gil et al., 2018; Stangl et al., 2018) noise within the path integrator might correspond to stochastic processes in the neurons or synapses of the grid cell system. Noise in the velocity input is likely to have a more diffuse origin in the sensing and sensory processing systems that extract velocity estimates from diverse sensory cues across the visual, vestibular, and proprioceptive pathways (Angelaki & Hess, 2005; Angelaki & Cullen, 2008; Laurens & Angelaki, 2011).

To refine our understanding of the source of noise in the integration pathway, we compared the default version of our model, in which the unbiased path integration noise accumulates with the travel distance along a trajectory, with a model variant with time-scaling of this noise. Direct comparisons between these two models showed that internal path integration noise mainly scales with traveled distance rather than elapsed time. This finding suggests that the main part of the accumulating noise in the path integrated location estimate stems not from noise intrinsic to the path integrator, which would tend to accumulate over time regardless of input, but from the sensing or sensory processing systems that compute self-motion estimates, and whose estimates must be noisy. Together with similar findings in non-spatial (Kiani et al., 2013) and spatial evidence accumulation tasks (Pinto et al., 2018), these results suggest an emerging principle in the neurobiology of integrators: that the dominant source of noise in the output of neural integrators is in the inputs rather than within the integrator circuit.

The present work also shows that path integration performance is reduced in older as compared to young adults, in line with previous studies (Mahmood et al., 2009; Adamo et al., 2012; Harris & Wolbers, 2012; Bates & Wolbers, 2014). Further, we were able to determine the dominant sources of error in older subjects and thus determine which of the sources of error already found in young subjects is most magnified as subjects age. Comparing the components of error in young and older adults revealed a significantly higher magnitude of unbiased noise in path integration computations of older adults, while other sources of error were not significantly different between age groups. In other words, the biggest source of error in young adults – accumulating unbiased noise likely arising from imperfect velocity estimation but possibly with some additional contributions of noise internal to the integrator – is further magnified in aging adults, while the smaller sources of error are not significantly compromised with age. Notably, older adults do not appear, at least in our experimental setup, to acquire major additional biases in their speed estimates or become substantially more inaccurate in their reporting of their internal location estimates. Rather, the most fragile part of the path integration process in younger subjects is most affected with aging.

What could potentially cause increased accumulating noise in older adults? Recent evidence from neuroimaging studies indicates that signatures of grid-cell-like activity in human entorhinal cortex are reduced in old age, and this reduction is associated with larger path integration errors (Stangl et al., 2018). Certain aspects of the vestibular system also deteriorate with age (for a review, see Allen et al., 2016), and vestibular loss affects path integration performance (Glasauer et al., 2002; Xie et al., 2017). These findings support the possibility that an increase in noise on a neuronal level, possibly from a loss of the signal-to-noise ratio from degrading circuitry in the grid cell or the vestibular system may be responsible for deficient computations of pose in old age. Alternatively, deficient processing of proprioceptive or optic flow information in cortical and sub-cortical structures could be responsible for noisy inputs to the path integration circuit in older adults. It therefore remains an important goal for future studies to determine which specific factors underlie increased internal noise in older adults’ path integration computations.

Together, we have shown here that path integration error in both young and older adults is mainly caused by accumulating unbiased noise, whereas other error sources contribute only modestly to total error. Moreover, we found that this noise is further magnified in older adults, and therefore accounts for the majority of age-related path integration deficits. Given the importance of path integration computations for spatial navigation, these findings not only advance our understanding of the specific contributors to path integration error, but may also shed light on the mechanisms that underlie navigational decline in old age.

## Methods

### Participants

62 healthy humans took part in this study. They had no reported history of neurological or psychiatric disease and no reported motor deficits during normal walking or standing. All participants reported right-handedness and had normal or corrected-to-normal eyesight.

Informed consent was obtained from all participants in writing before the measurements, and the experiment received approval from the Ethics Committee of the University of Magdeburg.

Prior to the study, all participants underwent the Montreal Cognitive Assessment (MoCA) screening tool for mild cognitive impairment (Nasreddine et al., 2005). Six older adults who did not exceed a MoCA cut-off score of 23 (following Luis et al., 2009) were excluded from the study and did not participate in any further measurements. Consequently, the data of the remaining 56 participants was used for data analyses: The group of young adults consisted of 30 participants (15 woman, 15 men) aged between 19 and 26 years (M = 22.0, SD = 2.0 years), whereas the group of older adults consisted of 26 participants (13 woman, 13 men) aged between 62 and 78 years (M = 69.0, SD = 4.6 years).

### Path integration task

Each participant’s path integration performance was measured using a behavioral path integration task, in which they had to track their own position during movement along pre-defined sinuous paths.

In commonly used path integration tasks for humans, such as the triangle completion task (Fujita et al., 1993; Harris & Wolbers, 2012), participants traverse a path and only estimate the distance and direction to the starting location at the end of the path. In the current study, we used a task in which participants were asked at four different points along the path to estimate the distance and direction to the path’s starting point (**Figure 1A**). Multiple distance and direction judgments per path were used for three reasons: First, it results in a larger number of data points (i.e., participant responses) in a similar amount of time, enabling a more reliable estimation of path integration errors. Second, it allows us to characterize the accumulation of the path integration error along longer and more complex paths. Third, responses from multiple points along the path can allow for a more precise estimation of path integration errors. Specifically, when complex paths are used, a participant may become disorientated in some trials as they move along the path, and the chances of this occurring increase with the distance traversed. When only one response is collected at the end of the path, as per the traditional triangle completion task, the participant’s estimate would be random and not provide a valid quantification of path integration performance. In contrast, our task samples from multiple points along the path meaning that, even if the participant has become disorientated at the path’s end point, there are still other data points earlier in the path that provide more accurate estimates of path integration performance.

Prior to the task, participants received written information about the task, and completed several practice paths. Participants donned a head-mounted display (HMD; Oculus Rift Development Kit 2, Oculus VR LLC, www.oculus.com), so that they could not see anything outside the HMD. During the task, participants wore earmuffs in order to prevent them from hearing any background sounds. Furthermore, they were instructed to immediately inform the experimenter if they noticed any external cues that could help them to orient during the task (such as hearing, seeing, feeling or smelling something).

During the task, participants held a wooden stick and were guided by the experimenter along a path (**Figure 1B**). At each of four stopping points along the path, the distance to the starting point had to be estimated verbally in meters and centimeters, and participants turned their body on the spot to indicate the orientation to the starting point, which was measured by the built-in gyrometer of the HMD.

Via the HMD, participants saw a virtual environment, which consisted of a ground plane and distant landmark cues (**Figure 1C**). The ground plane was designed to provide optic flow information during movement, but did not contain any fixed reference points or landmark cues. The distal landmarks were rendered at infinity, so that participants could use them only to determine their heading direction but not their position or any distance information. The exact position of a participant was tracked throughout the task using the Vicon Motion Tracking System with 12 cameras of type T10 (Vicon, Oxford, UK). The participant’s viewpoint within the virtual environment was constantly updated depending on their actual position and movement, so that participants could actively walk around in the virtual environment. Consequently, in order to keep track of their own position relative to the path’s starting point, participants could use both body-based and visual self-motion cues to perform the path integration task. Specifically, body-based self-motion cues included proprioceptive and vestibular representations, as well as motor efference copies that are produced during movement, whereas visual self-motion cues included optic flow information from the virtual environment and directional information from the environment’s distal landmarks (Etienne & Jeffery, 2004).

There were 10 different pre-defined paths (**Figure 1D**). Coordinates for each path were defined as follows: First, a 4-legged path was created that comprised four distances and three turning angles between them. Each distance was either 2, 3.5, 5, or 6.5 meters, and each angle was either 55, 80, or 105 degrees to the left or to the right. Various combinations of distances and angles were used, that fit into a rectangular area of approximately 10 × 8 meters (given by the tracking area and size of the room in which the experiment took place). On the basis of these 4-legged paths, we then created curved paths without corners by using the cscvn-function of MATLAB’s curve fitting toolbox to calculate a natural interpolated cubic spline curve passing through the turning points of the 4-legged path.

Six paths comprised a mixture of left and right turns, respectively (see **Figure 1D**, path numbers 1 to 6). Two additional paths (path numbers 7 and 9) only comprised right turns or left turns, respectively, and these two paths were present also in their mirrored version (i.e., the path that had only left turns was present also in its mirrored version comprising only right turns, and vice-versa). Directions (left vs. right) of the three turning angles per path were counter-balanced between the different paths.

Critically, the experimenters ensured that participants did not see the real physical dimensions of the testing room and the paths before and during the experiment, by guiding the participants into the room only after they had donned the HMD.

Participants completed the path integration task in three blocks. Within each block, participants performed each of the 10 paths one time and, in addition, they performed the paths 1 to 6 (the ones which had both left and right turns) another time without stopping at the first three stopping points but only at the end of the path (i.e., only at stopping point 4). Consequently, each participant performed 16 paths per block. The order of paths was pseudo-randomized, but the same order was used for all participants. There were always at least three different paths between repeats of the same path. The virtual environment was different in each block (see **Figure 1C**) and the order of environments was randomized across participants.

After the 4th and after the 12th path of each block, participants completed three so-called “standardization paths”, which were needed for data analysis in order to correct each participant’s distance estimate for their ability in verbally reporting distances using meter/centimeter units (see Methods section on “Calculation of path integration errors”). The procedure during a standardization path was similar as during a normal path, but a standardization path had only one start point and one stopping point, which were connected by a straight line, and participants had to estimate the distance between starting and stopping point. Three different distances had to be estimated in the following order: 10 meters, 2 meters, 6 meters. Moreover, there were short breaks in the middle of each block and between blocks. **Figure 1E** gives an overview over the procedure for each block.

After completing the task, participants filled out a form in which they were asked whether they noticed any external cues that could have helped them to orient during the task (such as hearing, seeing, feeling or smelling something), but no participant reported such confounding sources of information. Further, all participants were asked whether they had recognized that some paths were repetitions of each other, but no participant did.

The path integration task was developed using the WorldViz Vizard 5.1 Virtual Reality Software (WorldViz LLC, www.worldviz.com).

### Calculation of path integration errors

At every stopping point of a path, participants had to estimate the distance to the path’s starting point verbally in meters and centimeters. Converting an internal estimate of location to a verbal estimate is known to be biased (Izard & Dehaene, 2008). Here we assume that the bias is multiplicative. To measure the bias, we ask subjects to walk on straight standardization paths of length 2m, 6m and 10m and to report verbally the distance to the starting point. The correction factor for the bias is then given by:

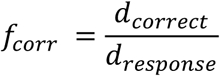

where *d_correct_* is the correct distance of the standardization path (2m, 6m, or 10m, respectively), *d_response_* is the responded distance, and *f_corr_* is the resulting correction factor. For each participant, this led to three different correction factors, one each for shorter (derived from the 2m standardization path), middle (derived from the 6m standardization path) and longer distances (derived from the 10m standardization path). These factors were used to standardize the distance estimates this participant reported at normal paths: Whenever the participant’s response distance of a normal path was between 0m and 4m, the response was multiplied with the correction factor for shorter distances, whereas response distances between 4m and 8m were multiplied with the correction factor for middle distances, and response distances larger than 8m were multiplied with the correction factor for longer distances.

This standardization procedure was done for each block-half separately, in order to ensure that standardization was performed using an up-to-date correction factor that also accounts for potential temporal changes of a participant’s perception of meter/centimeter units that might occur over the course of the experiment: Responses for the first half of each block (1st path to 8th path) were standardized using correction factors from the first set of standardization paths (i.e., carried out after the 4th path of a block), whereas responses for the second half of each block (9th path to 16th path) were standardized using correction factors from the second set of standardization paths (i.e., carried out after the 12th path of a block).

At each stopping point, the responded distance (multiplied with the respective correction factor *f_corr_*) and orientation was used to calculate the “presumed starting point”. The x and y coordinates of the presumed starting point according to the participant’s response were calculated by:

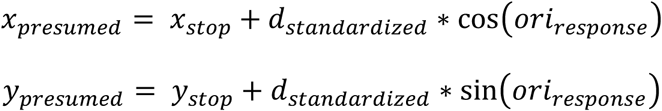

where *d_standardized_* is the standardized response distance, and *ori_response_* is the responded orientation. *x_stop_* and *y_stop_* are coordinates of the stopping point, *X_presumed_* and *y_presumed_* are the resulting coordinates of the presumed starting point.

To calculate the so-called “absolute” path integration error *Err_abs_*, we then calculated the Euclidean distance between the presumed starting point and the path’s correct starting point by:

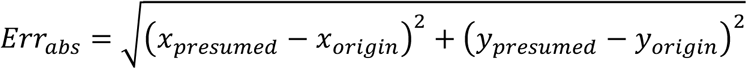

where *x_origin_* and *y_origin_* are the x and y coordinates of the path’s correct starting point. According to this method, each absolute path integration error reflects the error that occurred between the path’s starting point and the respective stopping point (i.e., at stopping point 1 it reflects the error between the starting point and stopping point 1; at stopping point 2 it reflects the error between the starting point and stopping point 2; and so on). Accumulation of this error measure (i.e., absolute path integration errors) across all available stopping points, however, would lead to an overrepresentation of errors that occurred on early path segments (because these errors would be included for both earlier and later stopping points).

In order to allow for accumulation of path integration errors across stopping points, we therefore also used an alternative method to calculate the so-called “incremental” path integration error *Err_inc_*. For a given stopping point, the Euclidean distance between the presumed starting point (according to the participant’s response at this respective stopping point) and the previously presumed starting point (according to the response at the previous stopping point) was calculated by:

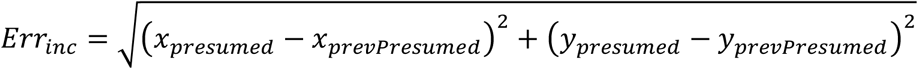

where *x_prevPresumed_* and *y_prevPresumed_* are the x and y coordinates of the previously presumed starting point (according to the response at the previous stopping point). Note that the previously presumed starting point at stopping point 1 is the correct starting point of the path (i.e., *x_prevPresumed_* = *x_origin_* and *y_prevPresumed_* = *y_origin_*). Consequently, this measure of the path integration error reflects only the incremental error that occurred on the latest path segment before the stopping point, but does not include the error that occurred on earlier segments of the same path. More specifically, at stopping point 1 it reflects the error that occurred between the starting point and stopping point 1, at stopping point 2 it reflects the error that occurred between stopping point 1 and stopping point 2 (not including the error between the starting point and stopping point 1), and so on. This method of calculating the path integration error allows, for each individual participant, to aggregate all error measures from all available stopping points, because each incremental path integration error measure includes only the incremental (i.e., unique) error contribution of one path segment.

### Computational modeling

The computational model we use differs from previous models of path integration error (e.g., Lappe et al., 2007, 2011) in several ways: First, we use time-resolved models in which moment-by-moment errors during a trajectory can interact with the moment-by-moment unfolding of the trajectory, and detailed, signed errors can be predicted over time. The richer model allows us to distinguish a large number of sources of noise and bias, and take into account reporting errors in which subjects are only able to report an imperfect representation of their internal location estimates. Unlike previous models that fit path-integration biases using trial-averaged data by minimizing the mean square error (Lappe et al., 2007, 2011), we model both biases and variances using a well-defined log-likelihood. This approach has several advantages: We can fit a more heterogeneous dataset where each trajectory is only repeated a few times, location estimates are weighted inversely proportional to the model-predicted variance (mainly influenced by the traveled distance), making the fit less biased and more data-efficient, and the log-likelihood allows a systematic model-comparison using cross-validation and the Bayesian Information Criterion (BIC).

Throughout this section we use bold-faced letters to refer to two-dimensional vectors. We assume that any participant continuously updates an internal, two-dimensional estimate 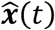 of their location ***x***(*t*) using an estimated walking velocity ***ν***(*t*). The update process is compromised by memory decay β, velocity gain α, additive bias **b**, and Gaussian noise **ξ**(t) with standard deviation σ_0_ (where **ξ**(t) is normally-distributed Gaussian noise) according to the following diffusion Langevin equation:

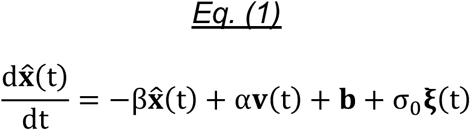

The parameters can be interpreted as follows:

- Memory decay or leak β: If β = 0, then 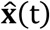 is the non-forgetful or perfect integral of the right-hand-side of the equation. If β > 0, then 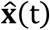 will have forgotten about inputs **v**(t − τ) with τ ≫ 1/β, thus the process is referred to as “leaky integration”.
- Multiplicative velocity gain or bias α: A value α > 1 corresponds to a systematic overestimation of displacement given velocity **v**(t), while a value α < 1 corresponds to an underestimate. Correct displacement estimation occurs when α = 1.
- Additive bias (AB) **b**: Specifies the bias direction along which the location estimate is pulled over time. Zero bias corresponds to **b** = 0.
- Accumulating noise (AN) that is unbiased and additive with standard deviation σ_0_: This noise can be interpreted to originate from a noisy integrator, a noisy velocity estimate input, or a mixture of both, depending on whether it adds up over time regardless of travel speed or if it scales with speed. Non-noisy velocity estimation and integration occur when σ_0_ = 0.

In our “full model”, we assume that the noise accumulates during displacements and thus grows in proportion to the travel distance. Therefore, the instantaneous value of σ_0_ is taken to be proportional to the square root of the instantaneous velocity magnitude (speed) |**v**(t)|. We consider variants in which this noise instead accumulates with elapsed time, independent of speed (see Methods section on “Model fitting, Extended Kalman Filter (EKF), and Bayesian model comparison” for more details). In a different variant, with constant noise (CN), noise does not accumulate at all but an overall unbiased Gaussian noise term whose total variance by the end of the trajectory does not scale with travel distance or time is added to the model estimate (see below). Within the accumulating noise models, the choice of an accumulating noise that scales with travel distance that we use in the full model, is better supported by our data (see Results section and **Figure 4A**).

Within the full model, we additionally assume that the subjects’ reports of estimated distance and angle to the starting point are corrupted by reporting noise (RN) (Schmidt et al., 1979; Jones et al., 2002; Faisal et al., 2008; Izard & Dehaene, 2008). Given an internally estimated distance d and angle φ, we assume that the reported distances 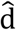 and angles 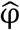 are given by:

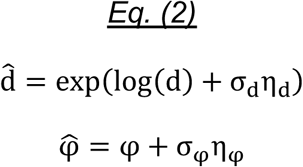

where σ_d_ and σ_φ_ are standard deviations of distance and angular noise, η_d_ is normally-distributed distance noise, and η_φ_ is normally-distributed angular noise. The parameterization of the distance reporting noise is chosen such that for fixed σ_d_, the magnitude of the reporting error 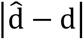 increases approximately linearly with d “proportional or Weber-like reporting noise” (RN), in line with Weber’s law (Oberlin, 1936; Gaydos, 1958; Cornsweet & Teller, 1965; Fechner, 1966; Indow & Stevens, 1966; Izard & Dehaene, 2008). We find empirically that this Weber’s law-type parameterization of the distance reporting error captures the data better than a linear parameterization, which we refer to as “constant reporting noise” (CRN) (see Results section and **Figure 3C**).

### Model fitting, Extended Kalman Filter (EKF), and Bayesian model comparison

Participants report their location estimates only at stopping points after moving along path segments. Before we can fit our model parameters to those estimates we first need to integrate the stochastic differential equation (1) along segments, a calculation that can be performed analytically because eq. (1) describes an Ornstein-Uhlenbeck process (Uhlenbeck & Ornstein, 1930; Pavliotis, 2014). Assuming that participants walk along a trajectory segment for time t with constant velocity **v**, the conditional distribution of the internal location estimate 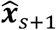 at the stopping point s + 1 given the estimate at the previous stopping point 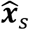 is given by the Gaussian distribution:

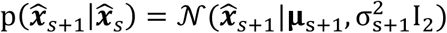

where I_2_ is the two-dimensional unity matrix and mean **μ**_s+1_ and variance 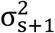 are given by:

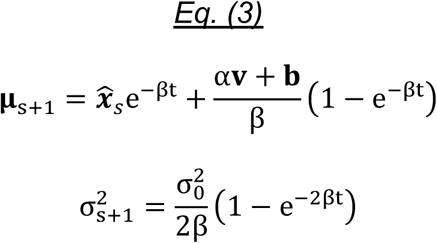

This update equation for the distribution of internal estimates can also be expressed in terms of the true length |Δ**x**| of the trajectory segment:

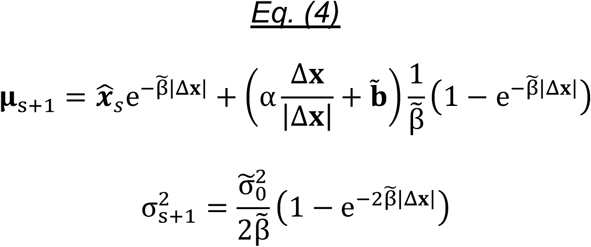

where we have rescaled three of the original parameters by the magnitude of the walking velocity |**v**|:

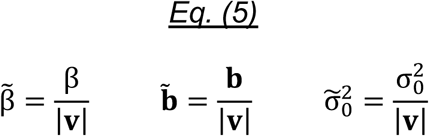

Equations (3) and (4) are equivalent if the walking velocity |**v**| is truly constant across trajectory segments and trials. If the walking velocity does vary, holding the transformed parameters (5) fixed assumes that the path integration error of the internal location estimate mainly depends on the traveled *distance*, whereas the original model (3) assumes that the path integration error mainly depends on the elapsed walking *time.* In what follows, we will choose the *distance* model and hold the transformed parameters (5) fixed, in line with previous modeling of human path integration (Lappe et al., 2007, 2011). We also explicitly test that the *distance* model is better supported by the data than the *time* model (see Results section and **Figure 4A**).

#### Fitting model without reporting noise (Full-RN)

Here we explain how the parameters 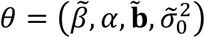 related to integration and *κ* = 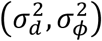 related to reporting were fit to participants’ performance by maximizing the likelihood. For simplicity, consider first a model without the reporting noise parameters *κ*. In this case the internal location estimate 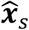 can be directly expressed in terms of participants’ report of the distance 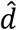 and angle 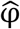 to the starting point **x**_start_ of the current walking trajectory:

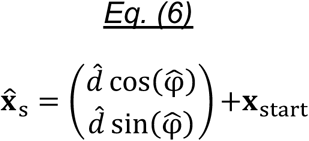

Without loss of generality we will set **x**_start_ = 0. The log-likelihood of the data averaged over trials is given by:

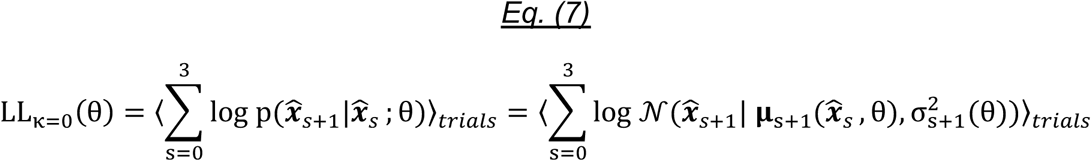

where 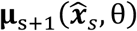 and 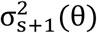 are given by the expressions in eq. (4). We then fit θ to the data by maximizing the log-likelihood numerically:

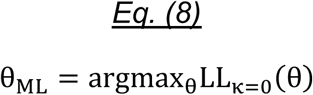

#### Fitting model with reporting noise: the full model (Full)

With reporting noise, the expression for the log-likelihood as a function of Θ = (θ, κ) is more involved, since the relationship between the reported estimates 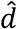 and 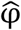 and the internal location estimate 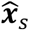 is both stochastic and non-linear. We can nevertheless make progress by rephrasing the problem in terms of the well-studied Extended Kalman Filter (EKF), a framework that permits calculation of the log-likelihood by locally linearizing the non-linearities (Thrun et al., 2005). The EKF framework encompasses a stochastic state transition of a hidden variable 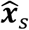 whose distribution can be inferred using a noisy observation z_s_:

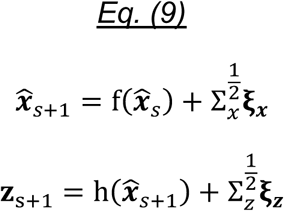

where f and h are arbitrary non-linear functions and Σ_x_ and Σ_z_ are covariance matrices of Gaussian-distributed noise. In our case the state transition is linear in 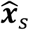 and is given as before by eq. (4):

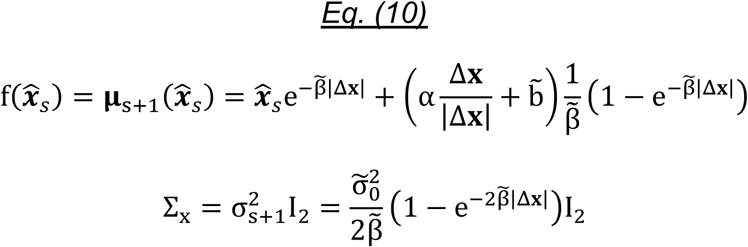

To derive the non-linear observation function we need to find a coordinate transformation such that in the transformed frame the noise is added linearly. According to eq. (2), the noise is added linearly in log-polar coordinates. The observation function 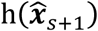 therefore corresponds to the transformation from cartesian to log-polar coordinates:

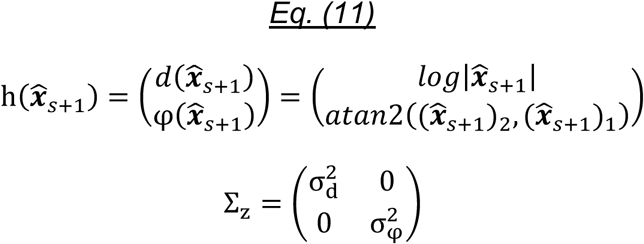

and the observation **z**_s+1_ is related to the reports 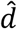 and 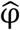 by:

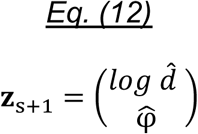

The EKF framework permits the calculation of two important distributions using Gaussian approximations: the posterior distribution of the hidden variable 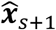 given the observations **z_1_** to **z**_s_ (predictive distribution), and the posterior distribution of 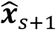 given **z**_1_ to **z**_s+1_ (updated distribution). We denote the mean and covariance of these posterior distributions as

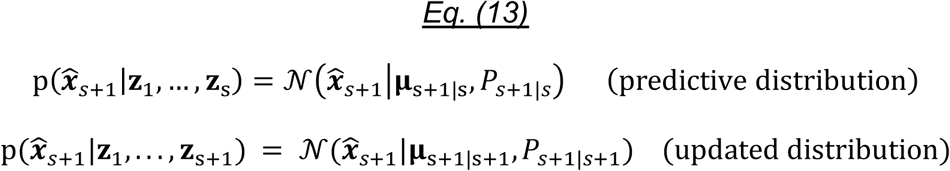

Mean and covariance of both distributions can be calculated recursively over stopping points using the standard EKF update equations (Thrun et al., 2005):

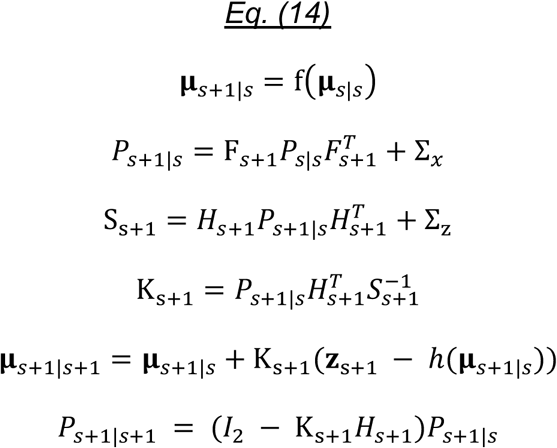

where the matrices F_*s*+1_ and *H*_*s*+1_ are the Jacobian matrices of transition and observation function evaluated at the previous updated mean **μ**_*s|s*_ and predictive mean **μ**_*s*+1|*s*_ respectively:

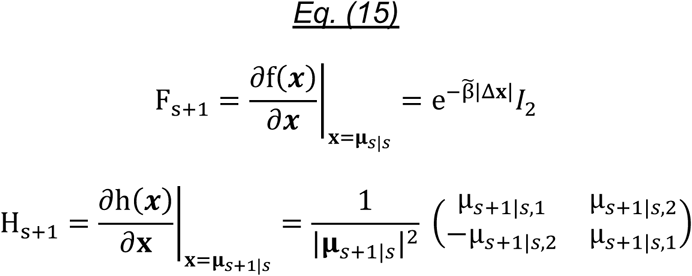

At the starting point (s = 0), we initialize **μ**_*s*=0|*s*=0_ = **x**_start_ = 0 and *P*_*s*=0|*s*=0_ = 0. Next, we calculate the predicted distribution of the next measurement **z**_s+1_ given the previous measurements **z**_1_ to **z**_s_ by integrating out the internal estimate 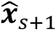:

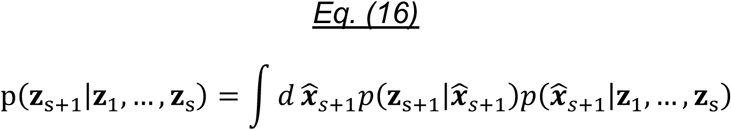

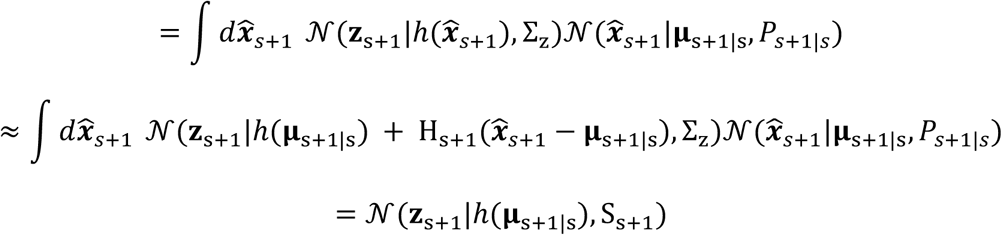

where we have used the linearization approximation of the EKF at the 3^rd^ line. This allows us to express the full log-likelihood as:

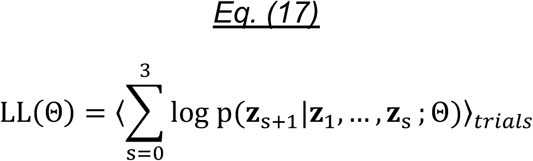

where the dependency on the parameters Θ is introduced through f, its Jacobian F_s+1_, Σ*_x_* and Σ_z_. In analogy to (8), we find the maximum likelihood (ML) estimate for Θ by numerically maximizing the log-likelihood:

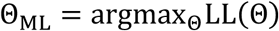

Numerical parameter optimization was performed using the fminunc-function of MATLAB’s optimization toolbox.

#### Incorporating trials without participant responses at intermediate stopping points

For a fraction of the trials, a response is not collected at intermediate stopping points, but only at the end of the trajectory. For these trials the observations **z**_s+1_ are missing for s ∈ {0,1,2} and therefore the EKF update equations (14) need to be adapted. This can be achieved using the infinite observation noise limit Σ_z_ → ∞, under which the predicted and updated posterior distributions become identical:

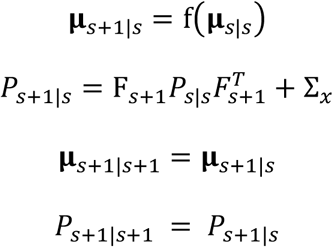

For s = 3, the observation at the last stopping point **z**_s+1_ is defined, and eq. (14) can be used as usual.

#### Model predictions

We simulated participants’ responses by sampling 100 repetitions of model trajectories for each participant and trial from eq. (9) given the fitted parameters Θ = Θ_ML_ and the trajectory parameters Δ**x** for each segment. Each repetition generates stochastic observations 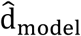 and 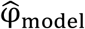 via eq. (12) that can be analyzed analogously to the actual data. The model prediction for the square error is calculated by averaging the square error of the simulated data over trials and repetitions. The model prediction for the bias on individual trials is calculated by averaging the simulated data over repetitions.

#### Model variants

##### Full model without additive bias, no reporting noise (Full-AB-RN)

The non-zero parameters in this model are memory decay 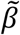, multiplicative velocity gain *α* and noise 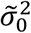. The additive bias (AB) 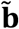 and reporting noise (RN) parameters *κ* = 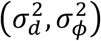 are set to zero. The log-likelihood is computed using eq. (7) instead of eq. (17).

##### Full model, no reporting noise (Full-RN)

This model has non-zero parameters 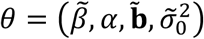 but the reporting noise parameters 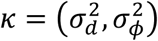 are set to zero. The log-likelihood is computed using eq. (7) instead of eq. (17).

##### Non-accumulating noise, no reporting error (Full-AN+CN-AB-RN, Full-AN+CN-RN)

These models assume that the total amount of noise is independent of distance, time, and stopping points, and the reporting noise parameters 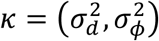 are set to zero. The fitting procedure for the non-noise (bias) parameters is equivalent to minimizing the square error in predicting the mean location estimates averaged over trials with similarly shaped trajectories. We replace Eq. (4) by:

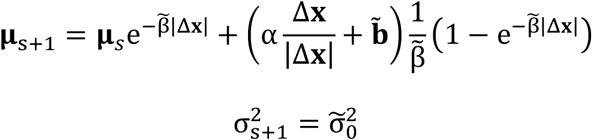

Note that **μ**_s+1_ depends on the previous predicted mean **μ**_s_ instead of the measured internal estimate 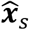 as in eq. (4). Correspondingly the conditional distribution of each internal location estimate does not depend on the estimate at the previous stopping point, so that 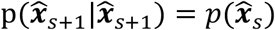. Maximizing the log-likelihood in eq. (7) corresponds to uniformly minimizing the square error across stopping points:

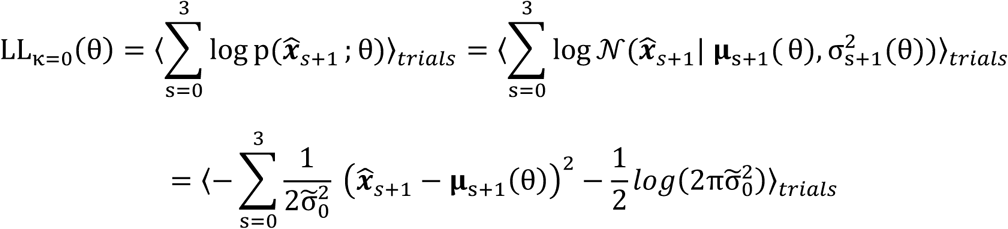

We fit two versions of the constant or non-accumulating noise model, one without any additive bias (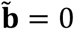; Full-AN+CN-AB-RN), and one with an additive bias (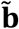 can vary; Full-AN+CN-RN). The model without additive bias (Full-AN+CN-AB-RN) is the closest match to the model proposed in Lappe et al. (2007, 2011), as it contains leak and bias.

##### Non-accumulating noise with reporting noise (Full-AN+CN)

As above, this variant assumes that the unbiased noise is independent rather than accumulating over time or distance, but does include reporting noise with non-zero reporting noise parameters 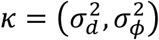, with Weber-like structure in which the reporting noise is proportional to the magnitude of the reported variable. This model can be fit using a few adjustments from the full model.

As there is no accumulating noise that induces correlations across stopping points, observations **z**_1_, …, **z**_s_ at previous stopping points are uninformative for the next location estimate 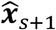, and both predictive and updated distribution in eq. (13) are equal to the prior distribution:

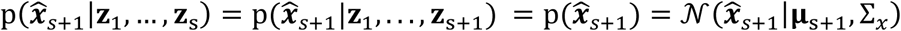

Consequently, there is no need to distinguish between predictive and updated mean and variance parameters. Instead, eq. (14) is replaced by:

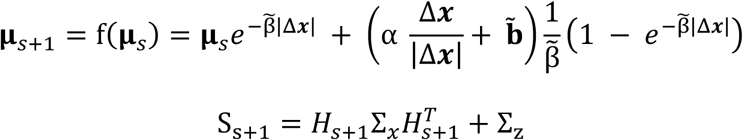

where

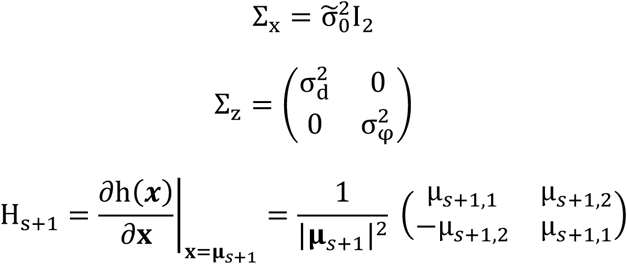

The log-likelihood is approximated as

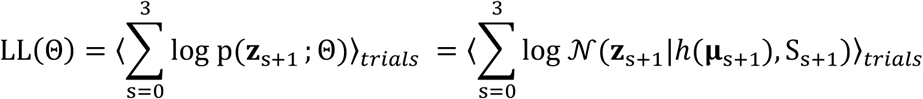

##### Model with constant reporting noise (Full-RN+CRN)

This model is the same as the full model (eq. (10)), except that the reporting error is drawn from a distribution of constant size, instead of being Weber-like (proportional to the reported quantity). Thus, eq. (2) is replaced by:

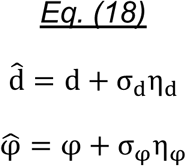

The model can be fit in the same way as the full model, when reporting noise is proportional to the internal estimate, except that noise is added linearly in polar coordinates instead of log-polar coordinates. Specifically, the first component of the observation **z**_s_ defined as the reported distance 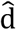 instead of its logarithm 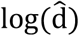, so that eq. (12) is replaced by:

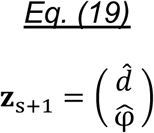

and we replace the observation function 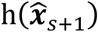 in eq. (11) by the transformation from cartesian to polar coordinates:

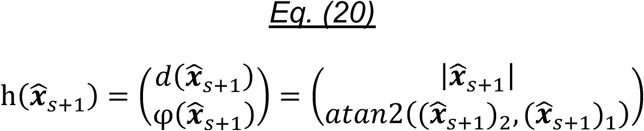

and the Jacobian H_s+1_ in eq. (15) by:

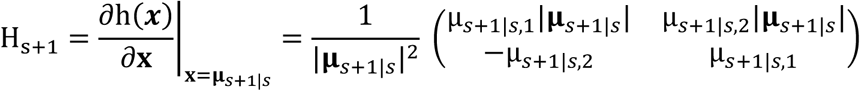

The rest of the calculation of the log-likelihood function is exactly the same as for the full model.

##### Fitting by age group

For this analysis, instead of fitting model parameters individually for each participant, participants in each age group are constrained to have the same model parameters.

##### Full model with time accumulation (ongoing noise is proportional to elapsed time rather than displacement; same reporting noise model as for the full model)

This model assumes that the mean and variance of the internal location estimate is determined by the elapsed time of each trajectory segment, eq. (3), instead of the distance of each trajectory segment, eq. (4). In the case of zero leak, the time model predicts that the variance of the internal location estimate increases proportionally to elapsed time instead of traveled distance.

To fit the time model we replace eq. (10) by:

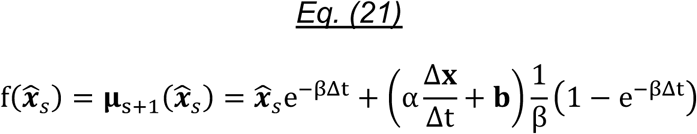

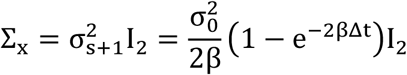

where Δt is the elapsed time of each trajectory segment. In addition, the Jacobian of the transition function F_s+1_ in eq. (15) is replaced by:

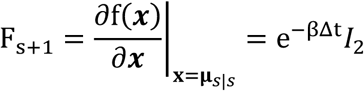

For trials without intermediate stopping points, only the total elapsed time of the trajectory, but not the elapsed time Δt of individual segments was recorded. For these trials we estimated Δt by linear interpolation using the traveled distance |Δ**x**| and assuming a constant walking speed.

The observation function 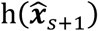 and its Jacobian H_s+1_ are identical to the standard Weber reporting noise model as specified in eq. (11) and eq. (15) respectively.

#### Model comparison using Bayesian Information Criterion and leave-one-out cross-validation

The Bayesian Information Criterion (BIC) is a scheme to compare models with different numbers of parameters: Models with lower BIC are preferred over models with higher BIC, and large BIC differences between models (ΔBIC ≫ 10) can be interpreted as “very strong” evidence against the model with lower BIC (Raftery, 1995; Konishi & Kitagawa, 2008). The BIC corrects for the higher expressibility of models with larger number of parameters using an additive compensation term. The formula for the BIC is given by:

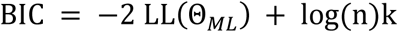

where n is the number of observations and k is the number of parameters. The number of parameters for different models is given by:

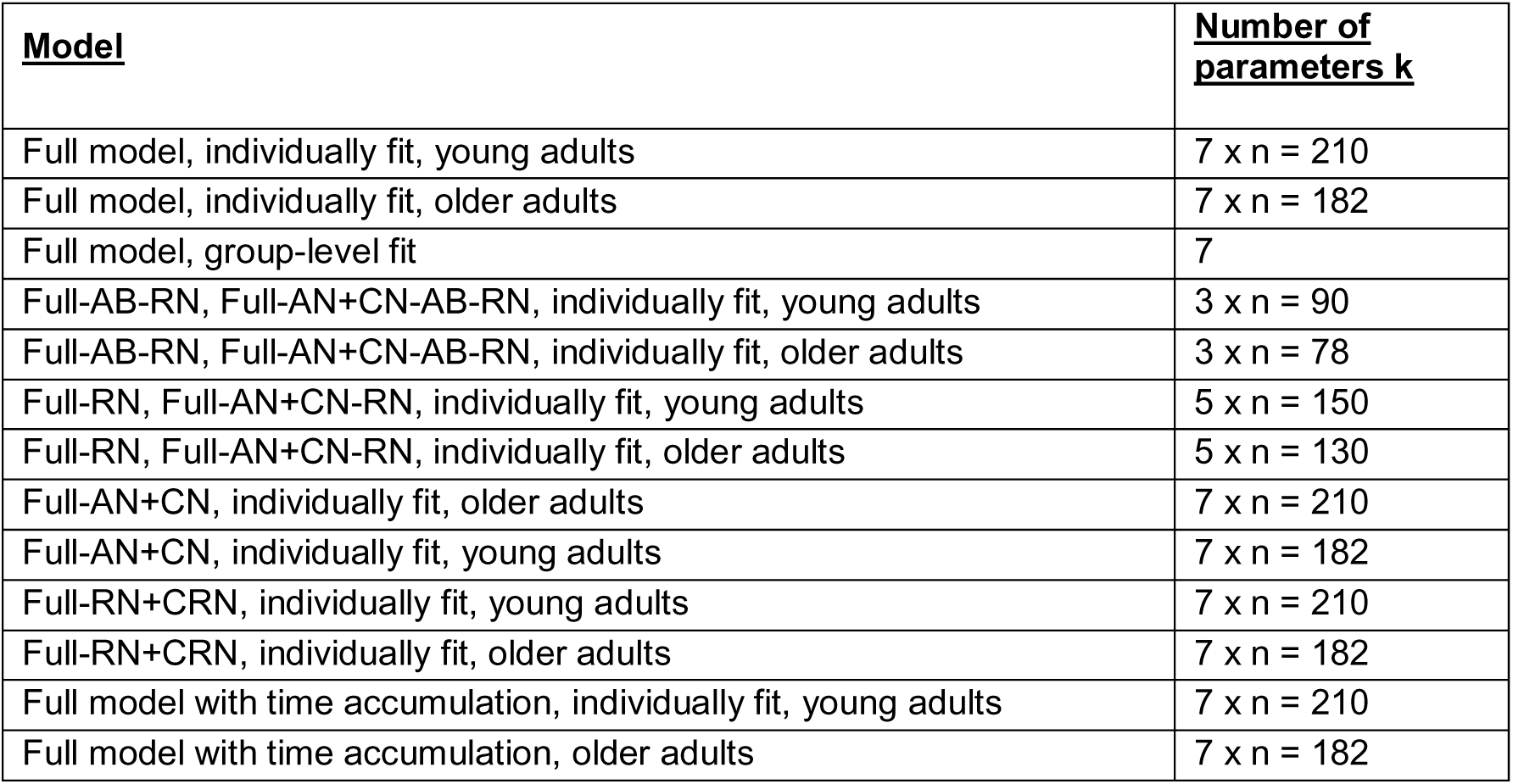

In addition to BIC, we compare models using leave-one-out cross-validation (LOOCV). Given T trajectories for each model and subject, we train the model parameters on a training dataset of T − 1 trajectories, evaluate it on the held-out test trajectory and average the result over the T distinct training-test splits. To allow numerical comparison with BIC we use as evaluation measure twice the negative log-likelihood:

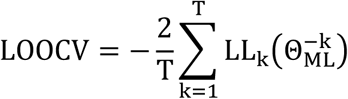

where LL_k_ is the log-likelihood corresponding to the k-th trajectory, and 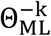 are the ML parameters on the training set excluding the k-th trajectory.

#### Relative influence of model parameters on predicted square error

The detailed computational model allows us to measure the influence of each type of bias and noise parameter on the square error predicted by the model. For each parameter type we calculated a reduced square error that is generated by setting this parameter type to its ideal value corresponding to unbiased, noiseless integration, while keeping the remaining parameters at their ML estimates:

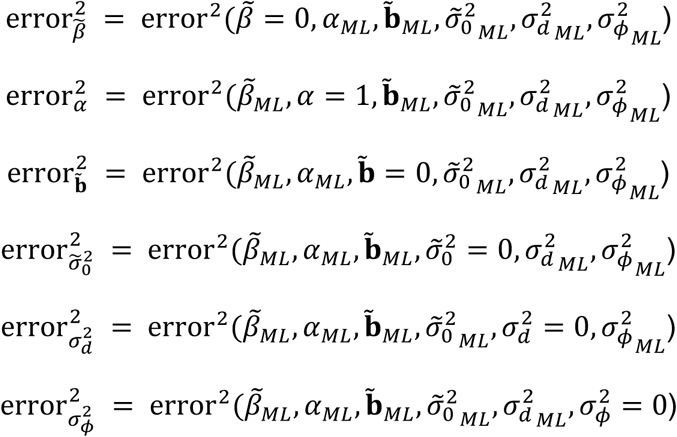

The relative influence of each reduced error in percent is then calculated as:

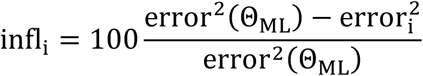

Note that the relative influence can be negative if the reduced square error is larger than the square error of the full model. This can be true in particular for the memory leak parameter 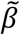: For example, a memory leak value 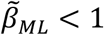 that draws location estimates towards the starting point can partly compensate for a velocity bias *α_ML_* > 1 that draws location estimates away from the starting point. Setting 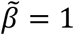 when *α_ML_* > 1 can therefore lead to a larger “reduced” square error and a negative relative influence.

Also note that due to the nonlinearity of the model, the relative influences do not have to sum to 100%.

## Author contributions

MS, IK, IF and TW conceptualized the work. MS and MR programmed and implemented the virtual reality path integration task. MS acquired the data. IK and IF developed the computational model for dissecting path integration errors. MS and IK analyzed the data, visualized the results, and drafted the manuscript. All authors edited the manuscript. IF and TW supervised the work.

## Acknowledgements

MS, MR, and TW express their gratitude to Falko Eckardt, Mareen Hanelt, Patrick Hauff, Anita Hökelmann, Marko Kirbach, Claudia Marx, Swantje Petersen, Mona Reißberg, and Uwe Sobieray for their help with data acquisition, project administration, and technical assistance. IK and IF thank members of the Fiete group for helpful discussions and comments.

This work was supported by a Collaborative Research in Computational Neuroscience Grant (01GQ1303) from the National Science Foundation (NSF) and the German Ministry of Education and Research (BMBF) to TW and IF; by the Simons Foundation through a SCGB grant to IF; and by the European Research Council Starting Investigator Grant AGESPACE (335090) to TW. IF is an HHMI Faculty Scholar and a CIFAR Senior Fellow.

---

**Supplementary Figure S1:**
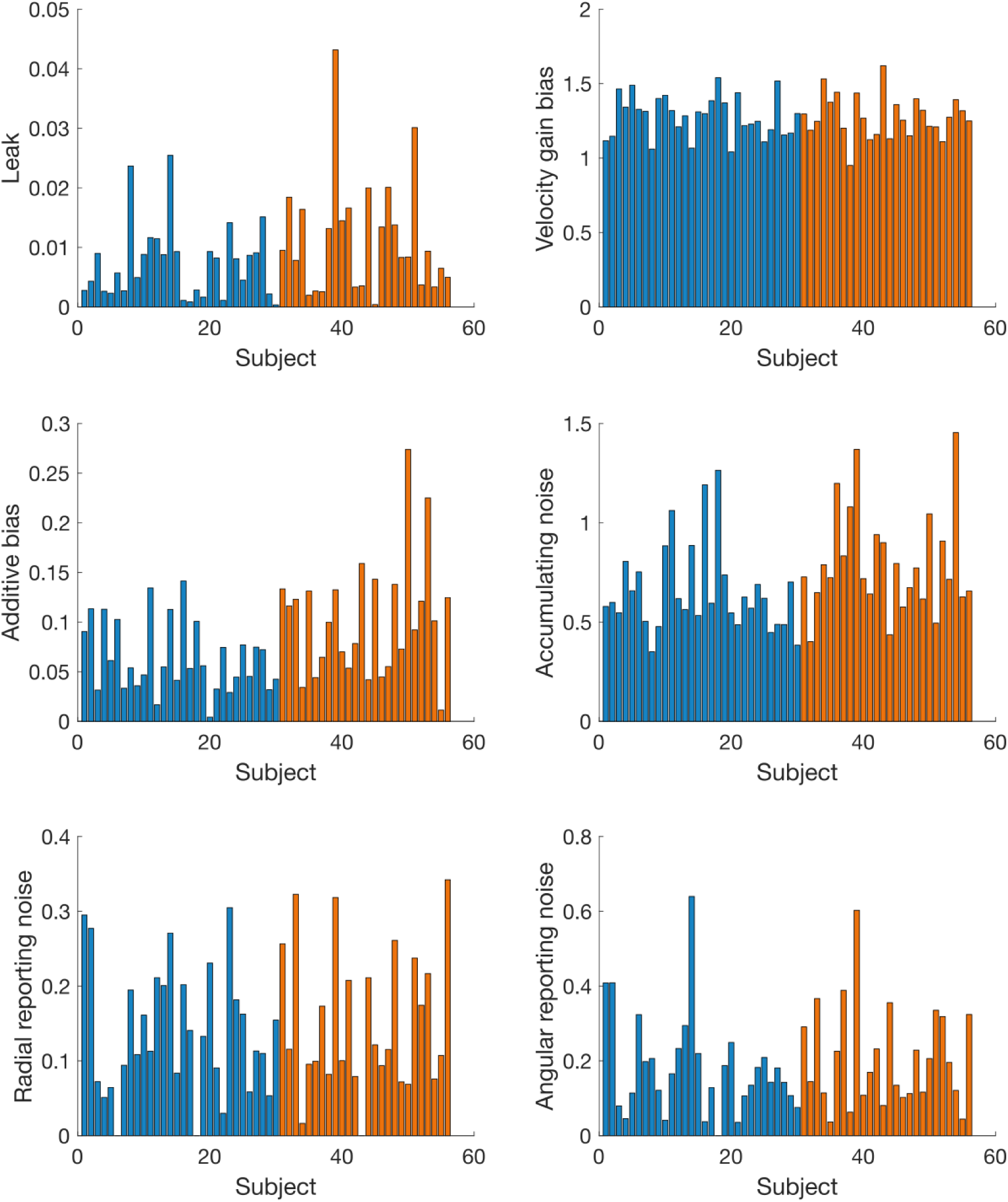
Individual parameter values for single subjects. Model parameter fit of the full model, shown for each subject individually. Blue bars indicate young subjects, orange bars indicate older subjects. Subject ordering is identical across plots.

**Supplementary Figure S2:**
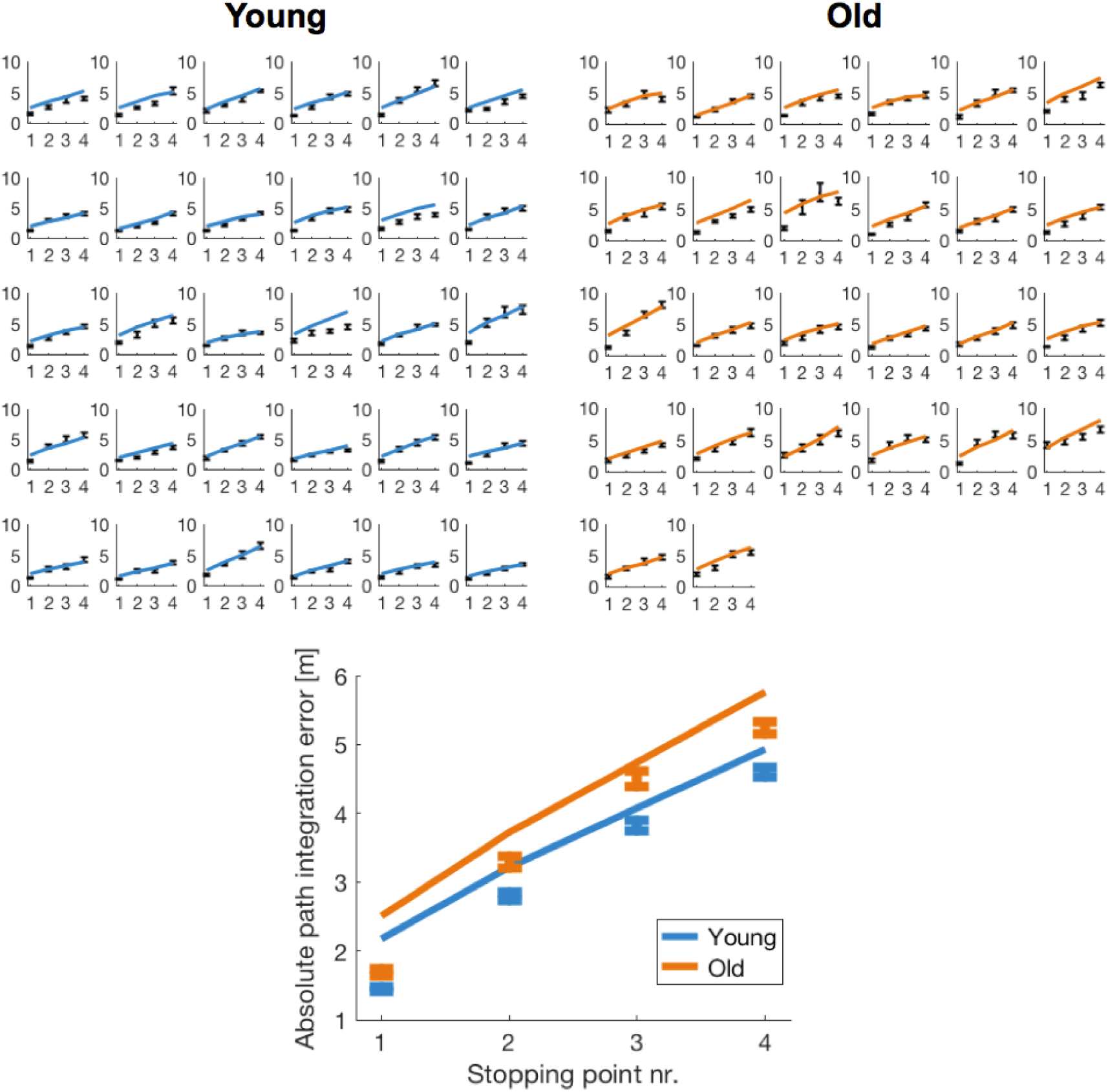
Path integration errors of young and older adults versus errors predicted by the model. **Top panel:** Path integration errors of each individual participant (black error bars) versus errors predicted by the model (solid lines) using each participant’s individual model parameters. **Bottom panel:** Mean path integration errors per age group (error bars) versus errors predicted by the computational model (solid lines). In order to calculate group-level model predictions, participants in each age group are constrained to have the same model parameters, instead of fitting model parameters individually for each participant. Error bars indicate mean ± SEM.

**Supplementary Figure S3:**
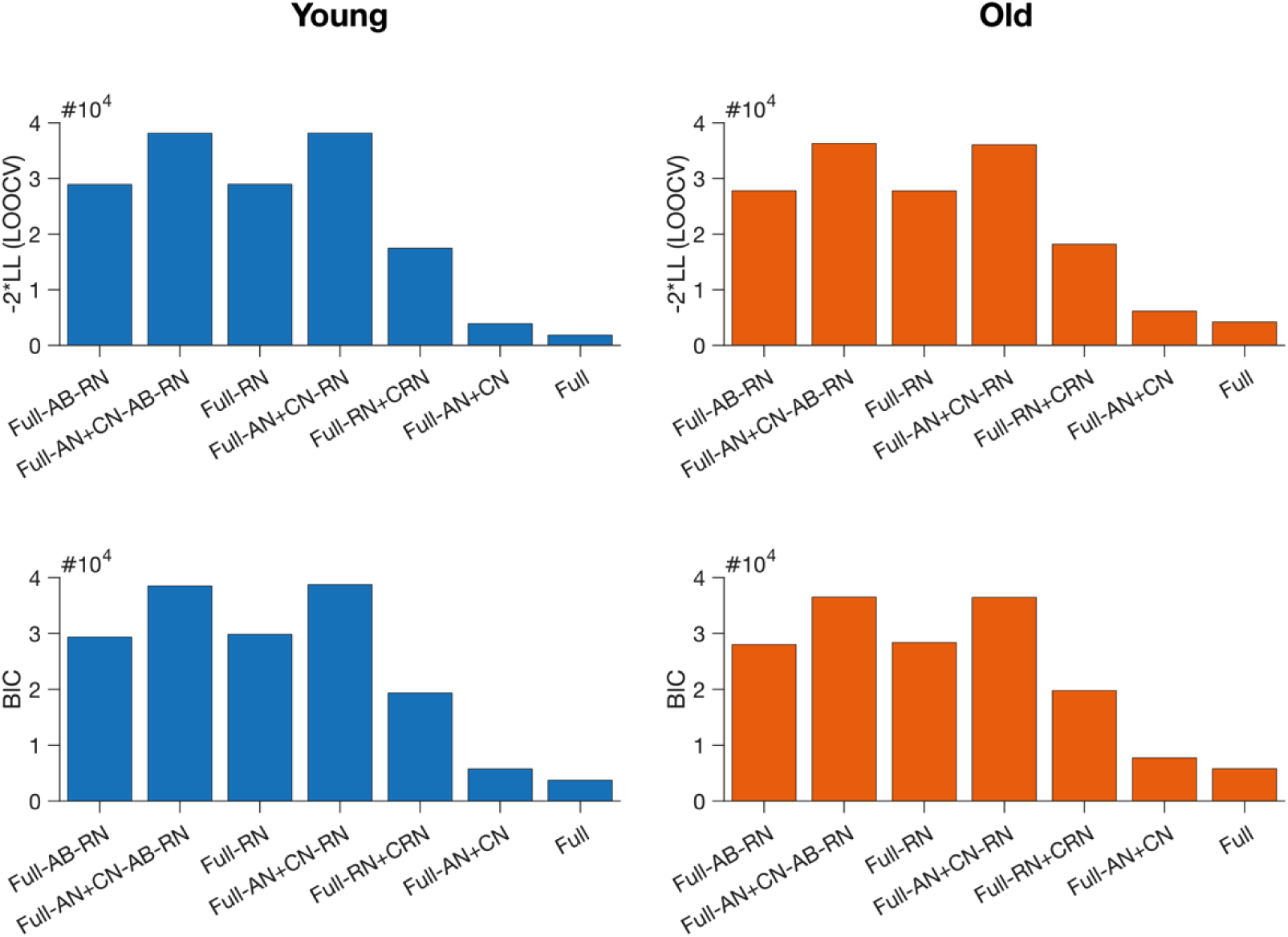
Comparison between model variants using BIC and LOOCV. Comparison between full model without additive bias, no reporting noise (Full-AB-RN); full model with constant instead of accumulating noise, no additive bias, no reporting noise (Full-AN+CN-AB-RN); full model without reporting noise (Full-RN); full model with constant instead of accumulating noise, no reporting noise (Full-AN+CN-RN); full model with constant reporting noise (Full-RN+CRN); full model with constant instead of accumulating noise (Full-AN+CN); and the default full modell (Full). For both age groups, the full model was best supported by the data. Higher bars indicate poorer model-fit. More details about different model variants and BIC/LOOCV model comparisons are provided in the Methods section.

**Supplementary Figure S4:**
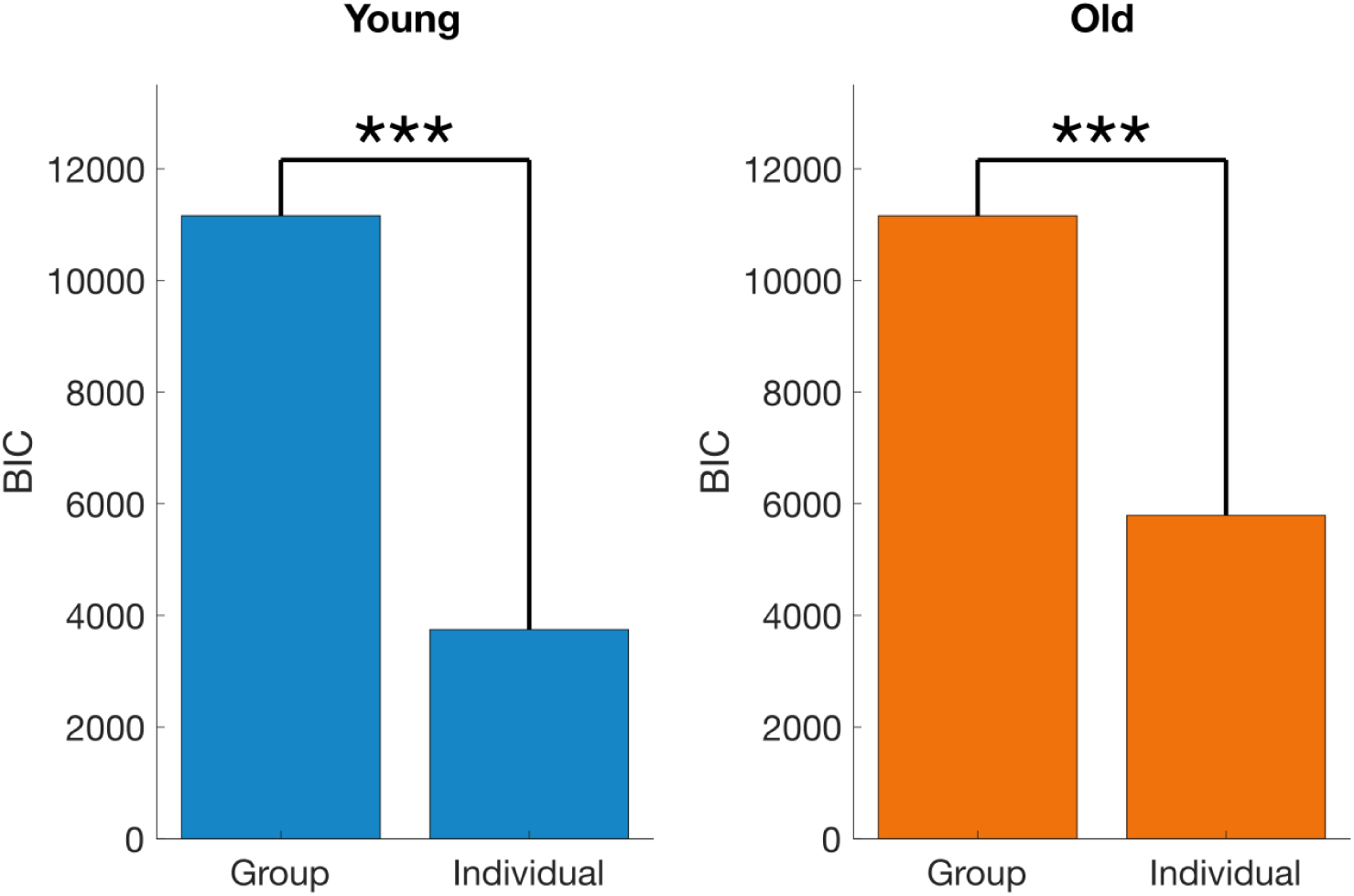
Comparison between group-level and individual models. Model comparison using BIC between models that were fitted at the group level and models that were fitted individually for each participant. For both age groups, the model with individual parameters per participant (i.e., the full model) was best supported by the data. Higher bars indicate lower model-fit. *** denotes “very strong” evidence against the model with lower model-fit (ΔBIC ≫ 10). More details about different model variants and BIC model comparisons are provided in the Methods section.

**Supplementary Figure S5:**
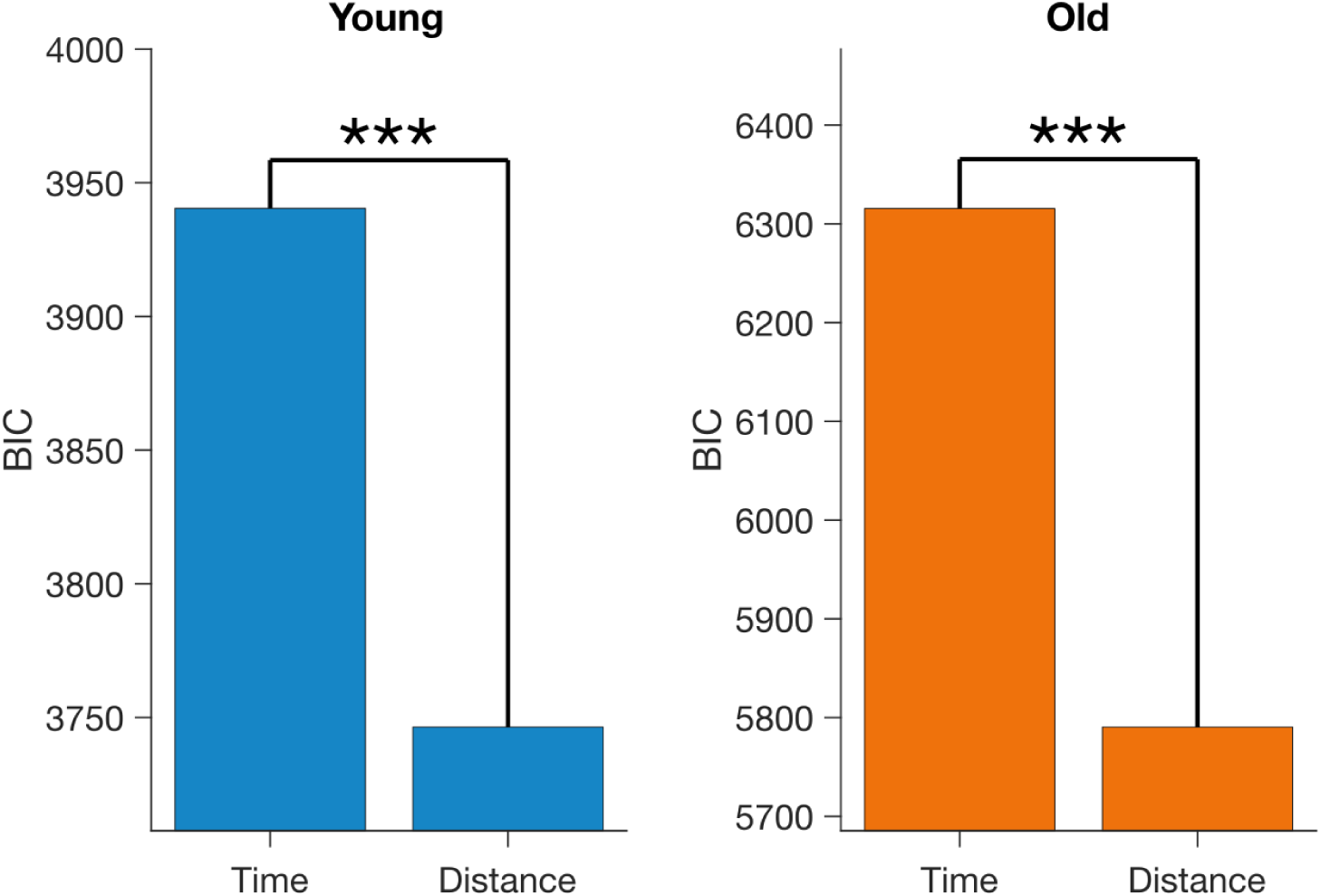
Comparison between “time model” and “distance model” variant using BIC. Model comparison using BIC between a model with time-scaling of the internal path integration error and the full model with distance-scaling. For both age groups, the full model is better supported by the data. Higher bars indicate lower model-fit. *** denotes “very strong” evidence against the model with lower model-fit (ΔBIC ≫ 10). More details about different model variants and BIC model comparisons are provided in the Methods section.

